# Several common methods of making vesicles (except an emulsion method) capture intended lipid ratios

**DOI:** 10.1101/2024.02.21.581444

**Authors:** Heidi M.J. Weakly, Kent J. Wilson, Gunnar J. Goetz, Emily L. Pruitt, Amy Li, Libin Xu, Sarah L. Keller

## Abstract

Researchers choose different methods of making giant unilamellar vesicles in order to satisfy different constraints of their experimental designs. A challenge of using a variety of methods is that each may produce vesicles of different lipid compositions, even if all vesicles are made from a common stock mixture. Here, we use mass spectrometry to investigate ratios of lipids in vesicles made by five common methods: electroformation on indium tin oxide slides, electroformation on platinum wires, gentle hydration, emulsion transfer, and extrusion. We made vesicles from either 5-component or binary mixtures of lipids chosen to span a wide range of physical properties: di(18:1)PC, di(16:0)PC, di(18:1)PG, di(12:0)PE, and cholesterol. For a mixture of all five of these lipids, ITO electroformation, Pt electroformation, gentle hydration, and extrusion methods result in only minor shifts (≤ 5 mol%) in lipid ratios of vesicles relative to a common stock solution. In contrast, emulsion transfer results in ∼80% less cholesterol than expected from the stock solution, which is counterbalanced by a surprising overabundance of saturated PC-lipid relative to all other phospholipids. Experiments using binary mixtures of some of the lipids largely support results from the 5-component mixture. Exact values of lipid ratios variations likely depend on the details of each method, so a broader conclusion is that experiments that increment lipid ratios in small steps will be highly sensitive to the method of lipid formation and to sample-to-sample variations, which are low (roughly ±2 mol% in the 5-component mixture and either scale proportionally with increasing mole fraction or remain low). Experiments that increment lipid ratios in larger steps or that seek to explain general trends or new phenomena will be less sensitive to the method used.

**SIGNIFICANCE STATEMENT:** Small changes to the amounts and types of lipids in membranes can drastically affect the membrane’s behavior. Unfortunately, it is unknown whether (or to what extent) different methods of making vesicles alter the ratios of lipids in membranes, even when identical stock solutions are used. This presents challenges for researchers when comparing data with colleagues who use different methods. Here, we measure ratios of lipid types in vesicle membranes produced by five methods. We assess each method’s reproducibility and compare resulting vesicle compositions across methods. In doing so, we provide a quantitative basis that the scientific community can use to estimate whether differences between their results can be simply attributed to differences between methods or to sample-to-sample variations.

## INTRODUCTION

Giant unilamellar vesicles (GUVs) come in many forms, from simple models of minimal membranes to complex mimics of living cells. Each application in which GUVs are used requires that different constraints are met, such as controlling vesicle size (1–3), number of lamellae (4–6), compositional asymmetry (7–9), membrane charge (10, 11) and encapsulation of solutes (12, 13). In response, many methods have been developed to produce GUVs (14), where each method prioritizes different constraints. However, when scientists use different methods to make GUVs, an uncomfortable question arises: do the vesicles produced by each method contain the same ratio of lipids as the stock solution from which the vesicles were made? If not, do all methods alter the intended lipid ratio in the same way, or does each method yield different ratios? These straightforward questions have huge impact: researchers who use different methods cannot compare their results unless they know that their vesicle compositions are similar or unless they know how to estimate the magnitude of the offset in lipid composition they might observe.

Concerns that different vesicle-making methods may incorporate different ratios of lipids into membranes are well-founded, especially when one of the lipids is a sterol. The solubilities of sterols in membranes of giant vesicles are sensitive to experimental conditions and to the identities of the other lipids in the membrane (15–18). Even when the mole fraction of cholesterol is below its membrane solubility limit, extrusion of vesicles may perturb the cholesterol fraction (19). Cholesterol levels are known to be especially low in vesicles made by emulsion transfer techniques (20–22). Challenges also arise in incorporating sufficient lipids with high melting temperatures (23), high charge (11, 24) or high spontaneous curvature (4) into membranes of giant vesicles.

Here, we used mass spectrometry to directly measure population averaged mole fractions of lipids in vesicles made by multiple methods. We chose lipids with a range of features that characterize broad classes of lipids (Fig. 1). Specifically, we chose cylindrical, zwitterionic lipids with both low and high melting temperatures (di(18:1)PC and di(16:0)PC), a charged lipid (di(18:1)PG), a cone-shaped lipid (di(12:0)PE), and a sterol (cholesterol). Each lipid was included for a different reason. PC-lipids produce stable membranes and are the most abundant lipid in mammalian cells (25–27). PG-lipids are abundant in mycobacteria (28). Their low pKa (29) and subsequent charge can increase the yield of vesicles made by gentle hydration, as do charged phosphatidylserine (PS) lipids (10). PE-lipids are one form of non-cylindrical lipids; PE-lipids facilitate membrane fusion and are abundant in the inner leaflet of red cell membranes (25, 27, 30). Cholesterol, in addition to facilitating membrane fusion (31), enables large-scale, liquid-liquid phase separation in membranes (32) and constitutes a large fraction (up to ∼40 mole%) of lipids in mammalian membranes (27).

**Figure 1.**
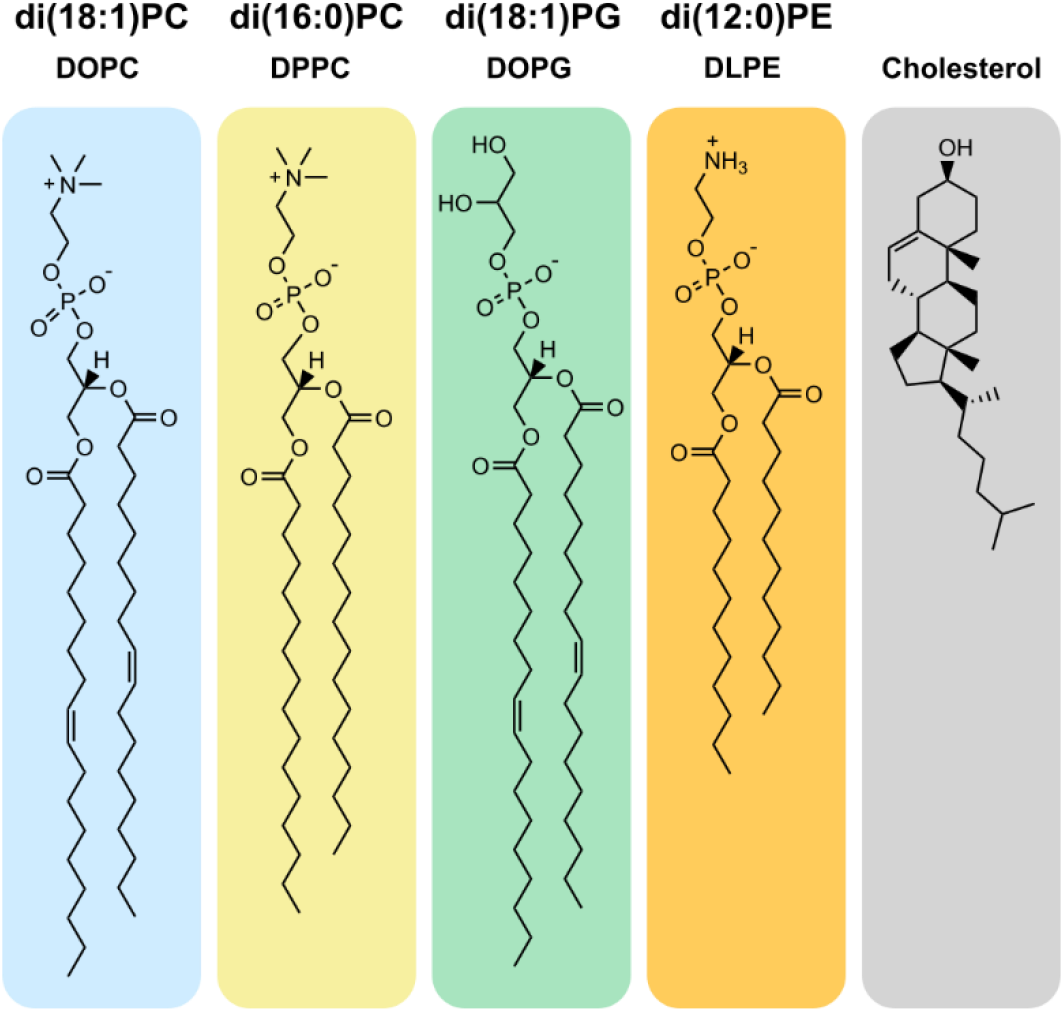
Structures of lipid types used to form GUVs. Lipids were chosen to represent five important lipid characteristics: saturated tails, unsaturated tails, charged headgroup, non-cylindrical shape, and sterol structure. Formal names of lipids reflect the length and unsaturation of the acyl chains and the headgroup type, whereas common names reflect historical sources. For example, di(18:1)PC (commonly called DOPC) is a zwitterionic lipid with a phosphatidylcholine (PC) headgroup and two 18-carbon chains, each with one double bond, whereas di(16:0)PC is saturated, with 16 carbons in each chain. Di(18:1)PG is a charged and unsaturated lipid with a phosphatidylglycerol (PG) headgroup. Phosphatidylethanolamine (PE) headgroups, as in di(12:0)PE, are smaller than PC headgroups, so PE-lipids are typically cone-shaped. Cholesterol has a hydroxyl moiety as a headgroup and a fused, four-ring structure with an 8-carbon chain.

We mixed these lipids in a 5-component mixture and in binary mixtures. We chose a single component mixture likely to produce high yields of vesicles by every method we used (Fig. 2). In detail, we chose our 5-component mixture to include high fractions of PC-lipids, to produce stabile membranes. We chose saturated (rather than unsaturated) PE-lipids, and we added them in low fractions, to avoid hexagonal phases and to minimize tubule formation (25). Similarly, we used low fractions of PG-lipids to enable electroformation. We ensured that fractions of cholesterol were well below its membrane solubility limit (15–18). To minimize vesicle-to-vesicle differences in composition, we chose all lipids to have melting temperatures ≤50°C (23). To answer key questions that arose from our experiments with the 5-component mixture, we then conducted experiments with membranes comprised of binary mixtures of lipids. Throughout, we compared vesicle compositions directly to stock solutions to minimize potential error due to variations in concentrations between different lots from lipid manufacturers.

**Figure 2.**
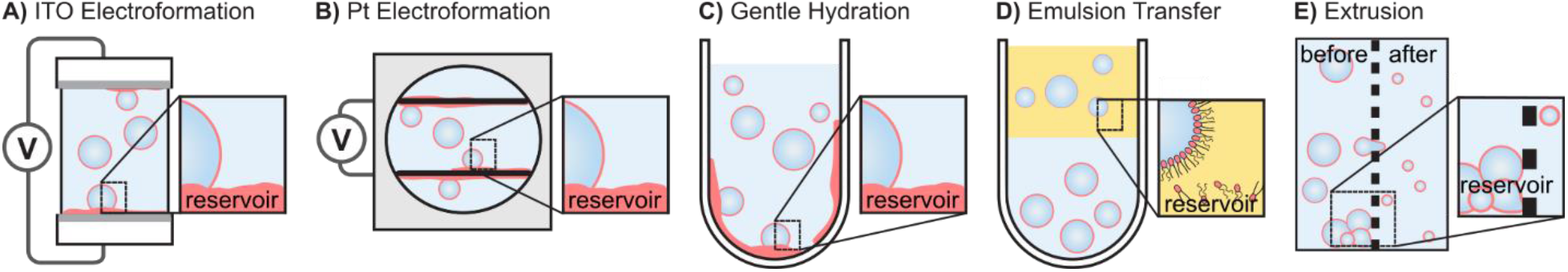
Methods of producing vesicles, showing reservoirs of unincorporated lipids. **A)** In “ITO electroformation”, an alternating field is applied across a hydrated lipid film on slides coated with indium tin oxide. Afterward, a residue of lipid remains on the slide. **B)** In “Pt electroformation”, an alternating field is applied across a hydrated film on platinum wires. Afterward, a residue of lipids remains on the wires. **C)** In gentle hydration, vesicles form spontaneously from a hydrated lipid film on a glass surface. Afterward, a residue of lipids remains on the glass. **D)** In emulsion transfer, lipids from an oil solution assemble at water/oil interfaces to form emulsion droplets, which then pass through a second interface to form vesicles. Afterward, a reservoir of lipids remains in the oil. **E)** In extrusion, smaller vesicles are made by passing giant vesicles through a porous filter. Afterward, a reservoir of lipids and vesicles remain on the filter. Figures are not to scale.

We chose four common techniques of producing GUVs: electroformation on slides coated with indium tin oxide (ITO), electroformation on platinum (Pt) wires, emulsion transfer, and gentle hydration (Fig. 2). We also included a common technique of converting giant vesicles into smaller vesicles: extrusion across a porous membrane filter. For each technique, we chose only one instance of experimental conditions commonly in use. Other labs may employ different conditions, so may observe different results.

Each technique for producing vesicles has advantages. Electroformation produces stable GUVs that span a wide range of sizes, frequently up to 100 µm (33). Emulsion transfer is valued for high encapsulation efficiency (34). Gentle hydration incorporates (and often requires) charged lipids in GUVs (10). Extrusion converts GUVs into vesicles of smaller and more uniform sizes. Each technique leaves a reservoir of excess lipids (Fig. 2, insets) that were not incorporated in the vesicles. For example, some common methods for gentle hydration of GUVs leave >50% of all lipids behind in the reservoir, and extrusion can incur a further loss of ∼20% of lipids (35, 36). Reservoirs may not contain the same ratio of lipids as the vesicles.

Our use of mass spectrometry to measure average vesicle compositions complements recent advances in using Time-of-Flight Secondary Ion Mass Spectrometry (ToF-SIMS) to directly measure vesicle-to-vesicle variations in lipid compositions (37). The two techniques are synergistic. Whereas current ToF-SIMS analyses excel at measuring relative compositions in individual vesicles, the technique currently has low sample-to-sample reproducibility for vesicle populations (37). Our mass spectrometry technique does the opposite. It excels in reproducibly evaluating average lipid ratios in vesicle populations and does not evaluate compositions of individual vesicles.

## MATERIALS AND METHODS

### Chemicals

PC-lipids, PG-lipids, PE-lipids, and phospholipid standards were obtained from Avanti Polar Lipids; cholesterol was obtained from Sigma-Aldrich; and d7-cholesterol standard was obtained from Kerafast. All lipids were used without further purification and without verifying concentrations because our experiments focus on measuring relative deviations in lipid composition from pre-mixed stock solutions rather than absolute concentrations in stocks. Lipid names reflect their headgroup (e.g., “PC”, “PG”, or “PE”), the number of carbon chains (e.g., “di” = 2), and each chain’s number of carbons and unsaturation (e.g., “18:1”). Specific lipids in our samples include dioleoyl-phosphocholine (di(18:1)PC or DOPC, *T*_melt_ = -18.3 ± 3.6°C (38)), dipalmitoyl-phosphocholine (di(16:0)PC or DPPC, *T*_melt_ = 41.3 ± 1.8°C (38)), 1,2-dipalmitoleoyl-sn-glycero-3-phosphocholine (di(16:1)PC or DPoPC, *T*_melt_ = -35.5°C (39)), dioleoyl-phosphoglycerol (di(18:1)PG or DOPG, *T*_melt_ = -18°C (40)), and dilauroyl-phosphoethanolamine (di(12:0)PE or DLPE, *T*_melt_ = 49.3 ± 1.7°C (41)). Lipid standards for mass spectrometry include di(15:0)PC, (16:0/18:1)PC, di(15:0)PE, di(15:0)PG, and d7-cholesterol. Cholesterol (Sigma-Aldrich) and a fluorescently labeled lipid (Texas Red DPPE, Life Technologies) were dissolved in chloroform at 10 mg/mL and 1 mg/mL, respectively. Mineral oil (light, bioreagent, 0.84 g/mL, 14.2-17.2 cSt at 40°C) was from Sigma Aldrich.

### Master Stocks

Seven master stock solutions of lipids (which included 0.8 mole% of the fluorescent dye Texas Red DPPE) were prepared in chloroform. Each stock was stored at -20°C in a vial sealed with Teflon tape, parafilm, and electrical tape. Lipid compositions of all master stocks were measured by HILIC-IM-MS as a baseline for direct comparisons between stocks and vesicle samples. The first of the seven stock solutions contained five lipid components (DOPC/DPPC/DOPG/DLPE/cholesterol). The second, third, and fourth contained binary mixtures that isolated the effects of unsaturation (DPPC/DPoPC), chain length (DPoPC/DOPC), and sterol content (DPoPC/cholesterol). The fifth, sixth, and seventh added 5 mol% DOPG to the binary mixtures, since charged lipids were required for the gentle hydration technique.

### Electroformation on ITO Slides

Electroformation followed standard procedures (33). Briefly, for each sample, 0.25 mg of total lipid (29 µL of master stock solution) was heated to 60°C and spread across two ITO-coated glass slides using the side of a glass Pasteur pipette. The slides were placed under vacuum for ∼30 min to allow residual chloroform to evaporate. An electroformation chamber was made by sandwiching 0.3 mm thick Teflon spacers between the ITO-coated slides. The chamber was filled with 300 mM sucrose (∼300 µL) and sealed with vacuum grease. The chamber was then attached to metal electrodes using stainless steel binder clips. An AC voltage of 1.0 V was applied across the electrodes at 10 Hz for 2 hr at 60°C. The temperature of 60°C is roughly 20°C above the highest lipid *T*_melt_ in the system. To minimize variability, all experiments using ITO slides were conducted by the same researcher.

### Electroformation on Platinum wires

A chamber was adapted from previous designs (42). Briefly, a vertical hole of 15 mm diameter was cut in a 25 mm x 25 mm x 5 mm block of Teflon. Two holes of ∼0.25-mm diameter were cut horizontally through the chamber, separated by 2.5 mm. Two 0.25-mm platinum wires were inserted into the two holes. Before each use, the wires and chamber were cleaned with chloroform. The bottom of the Teflon chamber was sealed with a glass cover slip, and the chamber was placed on a 60°C hot plate. Next, 5.7 µL of master stock solution was deposited in evenly spaced 0.5-µL drops on both wires. The interior of the chamber was filled with 1 mL of 300 mM sucrose solution, and the top was sealed with a glass cover slip. An AC voltage of 2.5 V was applied at 10 Hz for 2 hr at 60°C. To minimize variability, all experiments using Pt wires were conducted by the same researcher.

### Gentle Hydration

For each sample, 0.2 – 0.8 mg of total lipid from the master stock was transferred into a glass test tube. The test tube was placed in a water bath at > 50°C while chloroform was removed from the lipid solution by a steady stream of N_2_ gas to form a lipid film. To remove residual chloroform from the film, the test tube was placed under vacuum for ≥1 hr. After drying, the lipid film was rehydrated by adding 0.2 - 0.8 mL of 300 mM sucrose, so that the test tube contained 1 mL of solution for every 1 mg of lipid. The test tube was sealed with parafilm and maintained in an oven at 50°C for 24 hr. while vesicles formed. To minimize variability, all experiments using gentle hydration were conducted by the same researcher.

### Emulsion Transfer

The procedure was adapted from standard protocols (21, 34, 43) and was conducted at room temperature (20-25°C). First, a volume of master stock equivalent to 0.4-0.6 mg of total lipid mass was added to a chloroform-rinsed test tube. A lipid film was formed inside the test tube by drying the lipid solution under a gentle stream of nitrogen gas and placing the test tube in a vacuum chamber for ≥ 30 min. The test tube and a sealed container of mineral oil were quickly transferred to a glove box (Techni-Dome 360° Glove Box Chamber), which was then purged with nitrogen gas until humidity reached ≤ 25% (ThermPro Digital Indoor Thermometer). The container of mineral oil was opened only in the glove box, and 200-300 µL of mineral oil was added to the lipid film inside the test tube, resulting in a final lipid concentration of 2 mg/ml in mineral oil. The test tube was sealed with Teflon, parafilm, and electrical tape, removed from the glove box, and sonicated in a water bath (CO-Z Digital Ultrasonic Cleaner Model 10A, 40 kHz) for two or more intervals of 30 min at 50°C until no visible lipid film remained on the glass surface and the red color of the dye within the lipid-in-oil solution was homogenous. Between each 30 min sonication step, samples were gently vortexed to improve solubilization of the lipid film. The resulting lipid-in-oil solution was mixed with 20-30 µL (10% of the oil volume) of an aqueous, 300 mM sucrose solution by vigorous pipetting to create an emulsion. Approximately 200-300 µL of the emulsion was layered above 300 µL of an aqueous, 300 mM glucose solution in a 1.5 mL microcentrifuge tube and then centrifuged at 9000 g for 30 min to drive dense, sucrose emulsion droplets through the oil/water interface. The supernatant was removed without disturbing the vesicle pellet until minimal solution covered the pellet. The pellet was resuspended in 600-800 µL of 300 mM glucose by gentle pipetting, using tips that had been cut to larger diameters to minimize shearing. To minimize variability, all experiments using emulsion transfer were conducted by the same researcher.

### Vesicle Sedimentation and Storage

Vesicles made by electroformation and gentle hydration, vesicles were enriched in the solution (relative to lipid aggregates) by sedimenting vesicles in an osmotically matched glucose solution as in Fig. S1. Briefly, 200 µL of vesicle solution was added on top of 800 µL of glucose solution in a 13x1000 glass test tube (Fisher Scientific). Vesicles sank to the bottom of the tube for 10 min. Then 150 µL of vesicle solution was transferred from the bottom of the tube to a microcentrifuge tube. For emulsion transfer, vesicles were sedimented after resuspension in 600-800 µL of a glucose solution. For all methods, sedimented vesicle samples were stored at -20°C in parafilm-sealed 1.5 mL microcentrifuge tubes.

### Vesicle Imaging

Vesicle solutions were imaged between glass slides on a Nikon Eclipse ME600L upright epifluorescence microscope using a Hamamatsu C13440 camera at room temperature (25°C).

### Extrusion

Vesicle solutions were prepared by gentle hydration and enriched in solution (relative to vesicle aggregates) by sedimenting. The sedimented vesicle solution was split into two samples of equal volume. The first sample was reserved for “Before” extrusion measurements and the second sample was extruded to obtain an “After” measurement. To minimize loss of solution due to dead volume of the miniextruder (Avanti Polar Lipids), the filter supports and 0.1 µm polycarbonate filter were prewet by passing 1 mL of a sucrose solution through them prior to loading the sample. The complete extruder assembly (the heat block, the syringe with the vesicle solution, and the empty syringe) was maintained at 50 °C for 30 min before extrusion. Finally, the vesicle solution was passed through the filter 9 times and collected for analysis.

### Internal Standards

Cholesterol concentrations were normalized to the internal standard d_7_-cholesterol, which was dissolved in benzene at a concentration of 2.5 µg/mL. Phospholipids in 5-component samples were normalized to the following internal standards: both di(16:0)PC and di(18:1)PC were normalized to di(15:0)PC, di(12:0)PE was normalized to di(15:0)PE, and di(18:1)PG was normalized to di(15:0)PG. For binary samples, PC-lipids were normalized with respect to saturation: di(16:0)PC was normalized to di(15:0)PC, and both di(16:1)PC and di(18:1)PC were normalized to (16:0/18:1)PC. All phospholipid internal standards were 1 µM in samples.

### Lipid extraction

To enable quantification of each lipid class, internal phospholipid standards dissolved in chloroform and the d7-cholesterol internal standard dissolved in benzene were added to master stocks (in chloroform) and vesicle samples (in aqueous solution) before solutions were extracted for analysis. To maximize accuracy in comparing mass spectrometry data, sample concentrations were adjusted. Lipids were extracted from solutions by the method of Bligh and Dyer (44). Extracts were dried in a vacuum concentrator and reconstituted in 2:1 acetonitrile-methanol. Assuming each method of making vesicles incorporates 10% of the initial lipid mass into vesicles, vesicle solutions were diluted to achieve concentrations of ∼1 µM for the least prevalent lipid in the acetonitrile-methanol solution. Further details are in Fig. S17.

### HILIC-IM-MS Analysis of Phospholipids

Phospholipid analysis was conducted using hydrophilic interaction liquid chromatography (HILIC) coupled with ion mobility-mass spectrometry (IM-MS) in positive electrospray ionization mode (Waters Synapt XS HDMS; Waters Corp., Milford, MA). This technique relies on the presence of ionizable groups, which are present in lipids (and not in the hydrocarbons of mineral oil used in emulsion transfer techniques). Chromatographic separations were carried out using a Phenomenex Kinetex HILIC column (50 x 2.1 mm, 1.7 μm) with 95% acetonitrile/5% water/5 mM ammonium acetate as mobile phase A and 50% acetonitrile/50% water/5 mM ammonium acetate as mobile phase B (Waters Acquity FTN UPLC; Waters Corp.). Collisional cross-section calibration and IM-MS analysis were conducted as described previously (45–47). Data alignment and peak detection were performed in Progenesis QI (Nonlinear Dynamics; Waters Corp.). Retention time calibration and lipid identification were performed with the LiPydomics Python package (48). Lipid abundances were normalized to their internal standards and compared to the background. For only two types of samples (membranes produced by emulsion transfer in the main text that evaluate binary mixtures of lipids that differ in chain length and in PC-lipid vs. cholesterol), lipid abundances were deemed indistinguishable from the background. For these samples, extracts were concentrated by a factor of 5 and re-analyzed by HILIC-IM-MS.

### UHPLC-MS/MS Analysis of Cholesterol

Extracts were analyzed through ultra-high performance liquid chromatography-tandem mass spectrometry (UHPLC-MS/MS) using atmospheric pressure chemical ionization (Sciex QTRAP 6500; SCIEX, Framingham, MA or Waters Xevo TQ-XS; Waters Corp.) (49). Reversed-phase chromatography separations were carried out using a Phenomenex Kinetex C18 column (100 x 2.1 mm, 1.7 μm) with a 90% methanol/10% water/0.1% formic acid mobile phase. Quantitation methods and peak integration were performed using Analyst software (SCIEX Analyst 5.1; SCIEX) or MassLynx and TargetLynx software (Waters Corp.). Concentrations of cholesterol were obtained using peak ratios relative to d_7_-cholesterol (49) and a calculated average relative response factor (RRF) (for further information on RRF, see Tables S12, S16, and S18). Cholesterol abundances in emulsion transfer samples that evaluate binary mixtures of a PC-lipid and cholesterol were measured after concentration by a factor of 5.

### Analysis and Plotting

Phospholipid abundances are raw values obtained from the mass spectrometry measurements. To calculate lipid composition as mole percentages, the abundance of each lipid (normalized to a lipid standard) was divided by the total lipid abundance for each sample (Tables S11-20). Cholesterol abundances are raw values divided by the experimental RRF. The mean percentage of each lipid type and the standard deviation were calculated for three independent samples. Ternary plots were generated using the open-source Python library, python-ternary (https://github.com/marcharper/python-ternary).

## RESULTS AND DISCUSSION

Our experiments answer four types of questions: 1) Do all methods produce giant vesicles with sufficient yield? 2) How do the ratios of lipids in vesicles produced by each method differ from ratios in vesicles produced by the other methods, 3) How do those ratios differ from the stock solution, and 4) How reproducible is each method? We begin with a 5-component mixture of lipids and then conduct targeted experiments in binary mixtures.

### All methods produce sufficient vesicles from the 5-component mixture

As expected, all five methods produce vesicles. Vesicles produced by electroformation (on ITO slides and platinum wires) are predominantly unilamellar (Fig. S2 and S3) and are at sufficient yield for mass spectrometry. Vesicles produced by electroformation on Pt wires were smaller and less numerous than vesicles from electroformation on ITO slides (Fig. S3), which is consistent with fewer moles of lipids being deposited on Pt wires than on ITO slides. Vesicles made by gentle hydration consistently have more lamellae or are nested (Fig. S4), and vesicles made by emulsion transfer show membrane defects or nested vesicles (Fig. S5). Vesicles extruded through 100 nm membrane filters are too small to be resolved by traditional fluorescence microscopy (Fig. S6). After vesicles are formed by all techniques, lipids remain in the reservoirs shown in Fig. 2, either as residues on surfaces or dissolved in oil (Fig. S7-S10).

Vesicles made by electroformation and gentle hydration exhibit liquid-liquid phase separation of their membranes, consistent with the tendency for membranes to demix when they contain mixtures of high-*T*_melt_ lipids, low-*T*_melt_ lipids, and cholesterol (23). In contrast, the vesicles prepared by emulsion transfer did not exhibit liquid-liquid phase separation at room temperature, indicating that the lipid composition of emulsion transfer vesicles differed from the composition of the other vesicles (Fig. S5).

### Electroformation and gentle hydration create vesicles with minor offsets in lipid ratios

Our central result is that three methods of producing giant vesicles (electroformation on ITO slides, electroformation on platinum wires, and gentle hydration) result in similar population-averaged percentages of lipids in the membranes. Moreover, these percentages are similar to that of the master stock solution. These similarities are shown in the first four stacked bars in Fig. 3A.

**Figure 3.**
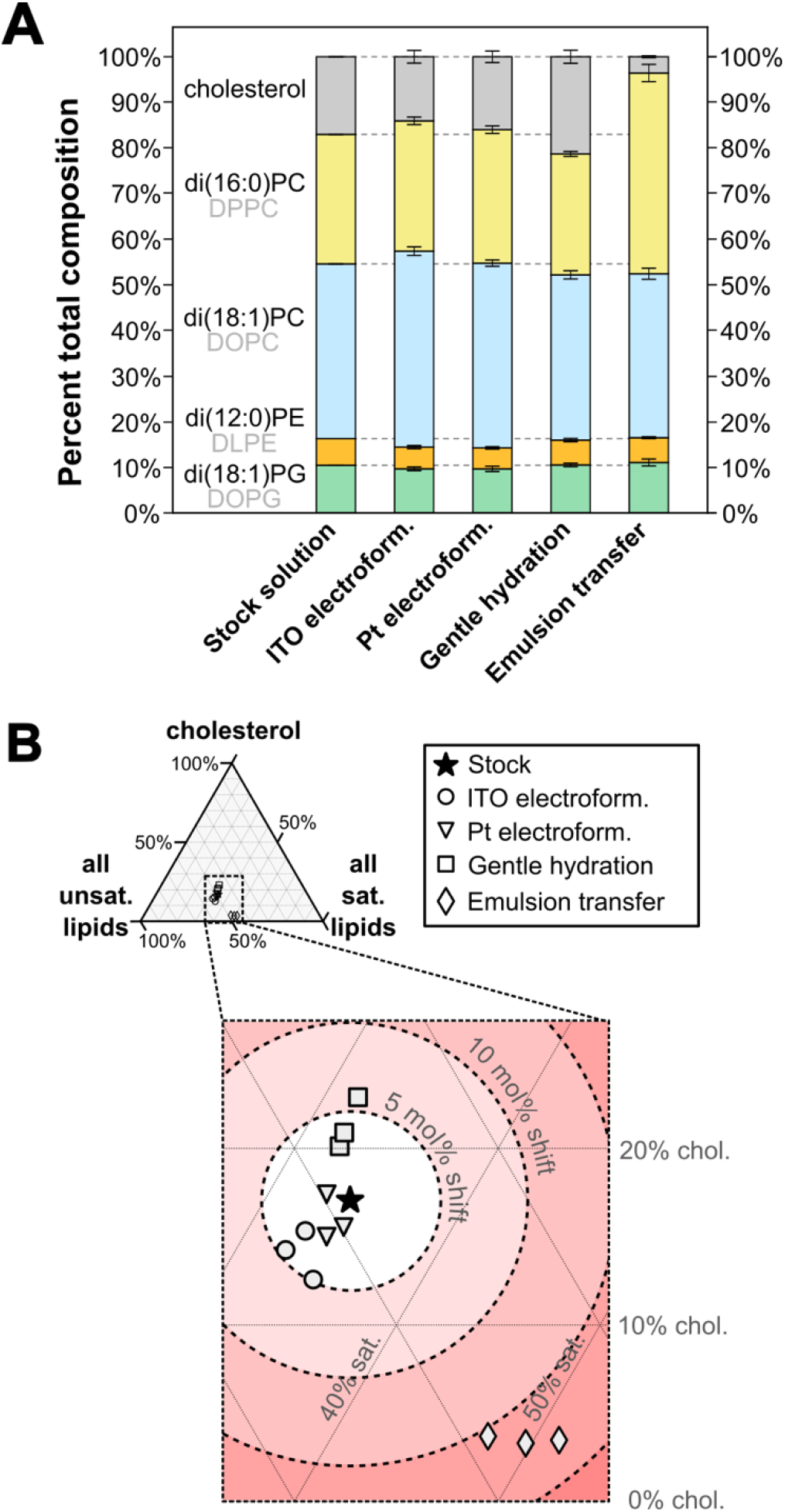
Lipid percentages in 5-component vesicles produced by different methods. **A)** Percent of each lipid in a 5-component stock solution and averages from three independent vesicle solutions made from the stock solution by each technique. Error bars above each section of the bar chart are standard deviations of the three independent experiments. Full data appear in Tables S1, S11, and S12. **B)** Lipid percentages for each independent experiment plotted on a pseudo-ternary diagram, where the three vertices represent cholesterol, the sum of the saturated lipids (di(16:0)PC and di(12:0)PE), and the sum of the unsaturated lipids (di(18:1)PC and di(18:1)PG).

Another way to visualize the minor differences in lipid ratios between the three methods is to zoom into a region of the pseudo-ternary diagram in Fig. 3B. Symbols for the three independent experiments of each method (circles for ITO electroformation, triangles for Pt electroformation, and squares for gentle hydration) cluster around the single star denoting the master stock solution. Fig. 3B shows that electroforming vesicles on ITO slides and on platinum wires shifts lipid mole ratios from the master stock solution by roughly 4 mole% and 1.5 mole%, on average. Similarly, gentle hydration of vesicles shifts lipid mole ratios by roughly 5 mole% on average, in the direction of increasing cholesterol. When expressed as a percent change, the gentle hydration method incorporated ∼20% more than the expected amount of cholesterol in the 5-component sample.

The data in Fig. 3 also show that reproducibility of these three methods is high, based on the close clustering of data from three independent experiments: the range of sample-to-sample lipid ratios for each method is within ±2 mole%.

Our result that electroformation and gentle hydration reproducibly create vesicles with similar mole fractions of cholesterol is consistent with reports that the maximum amount of cholesterol that can be incorporated into membranes is roughly equivalent for vesicles made by electroformation (between 65 and 70 mole% (18)), gentle hydration (61 mole% (50) or ≤ 63 mole% (15)), and a third method, rapid solvent exchange (66 ± 1 mole% (15)). Recent ToF-SIMS data disagree, but have low reproducibility: between two independent experiments, the amount of cholesterol in GUVs made by electroformation was either ∼1/4 or ∼2/3 of that in GUVs made by gentle hydration (37).

Shifts in lipid compositions in Figure 3 are largely due to different mole fractions of cholesterol in vesicles with respect to the master stock solution. In contrast, only small shifts of ∼1 mole% occur in PE-lipid and PG-lipids (Fig. S11). These shifts are the same magnitude as sample-to-sample differences of independent experiments. The small shift in di(12:0)PE seems surprising in light of previous studies using lipid mixtures extracted from rabbit sarcoplasmic reticulum: PE-lipids were grossly over-represented in the lipids left behind on a glass substrate (4). The particular lipid we use, di(12:0)PE, may integrate better into vesicles than other PE lipids because it is in a liquid, lamellar phase (rather than a gel phase or an inverted hexagonal phase). In contrast, the small shift in the charged PG-lipid is not surprising, at least with respect to gentle hydration of samples with high fractions of PG-lipids. Blosser et al. found agreement between the mole fraction of diphytanoylPG lipids in a stock solution and in vesicles that were made by gentle hydration, for molar ratios of >66% (11). Similarly, for a lipid mixture with ≥4% charged PS-lipids, Angelova found no significant difference in the mole fractions of the charged and uncharged lipids between a vesicle solution made by gentle hydration and a lipid film left behind on a glass surface (4).

How will the minor shifts in lipid ratios we observe in Fig. 3 impact how the biophysics community interprets data from GUVs? It depends on how experiments are designed. Broad conclusions derived from experiments that increment stock lipid compositions by ∼5 mole% will likely apply equally well to vesicles electroformed on ITO slides or on platinum electrodes, and may slightly differ when applied to vesicles made by gentle hydration. In contrast, detailed conclusions that depend on incrementing lipid compositions by only ∼1 mole% will likely apply only to vesicles made by a single method and will likely be affected by sample-to-sample differences. More detailed conclusions are difficult to extract from the data in Fig. 3 because only the percentages of the lipids (rather than their absolute values) are relevant. For example, an increase in the ratio of cholesterol to phospholipids could be due to an absolute increase in cholesterol, an absolute decrease in phospholipids, or both. Similarly, it is difficult to speculate whether our results would hold for every implementation of these techniques (e.g., with different solutions).

### Emulsion transfer results in too little cholesterol in vesicles

The emulsion transfer method we used creates large shifts in lipid compositions of vesicles made from 5-component lipd mixtures (Fig. 3). The largest of these shifts is that the emulsion transfer method incorporated 80% less cholesterol than expected from the master stock solution. This result is consistent with previous reports of severe reduction in cholesterol for vesicle membranes made by emulsion techniques such as cDICE (<1% of the expected cholesterol (20), double-layer cDICE (25-35% of the expected cholesterol (22)), and emulsion phase transfer (28-50% of the expected cholesterol (21)), as discussed in (20)). The problem likely arises because cholesterol lacks a large polar headgroup to drive it out of bulk oil, toward an interface with water. The problem can be mitigated, but not eliminated, by switching from heavy (20) to light mineral oil or by adding higher levels of cholesterol to the oil than the target level desired in membranes (21). Given that the choice of oil impacts cholesterol incorporation into vesicles, the exact values we measure apply only to one way of implementing the technique.

### Emulsion transfer exhibits large, chain-dependent shifts in PC-lipids

The emulsion transfer method we used vastly skews the ratio of the PC-lipids in 5-component lipid mixtures. Specifically, (16:0)PC is over-represented by a factor of ∼1.5 in the vesicles compared to the master stock solution (Fig. 3), an absolute increase from ∼30% to ∼45%. The effect is so large that nearly all loss of cholesterol in emulsion transfer vesicles appears to be counteracted by a gain in one of the phospholipids, di(16:0)PC. For comparison, the other PC-lipid, di(18:1)PC, decreases by only ∼2 mol%. To our knowledge, shifts in phospholipid compositions this large have not previously been measured in emulsion transfer vesicles. Shifts in the relative amounts of PC-lipids in vesicles made by other techniques (ITO electroformation, Pt wire electroformation, and gentle hydration) are minor. The largest of these is an increase of ∼5 mol% for di(18:1)PC by ITO formation (Fig. 3A). Other shifts in PC-lipids are on the order of 2 mol%, consistent with the scatter from independent trials in Fig. 3B. Similarly, shifts in PE-lipid and PG-lipid are on the order of sample-to-sample variations (Fig. S11).

### Binary vesicles show shifts in lipid ratios due to unsaturation and cholesterol content

Two structural characteristics of PC lipids, their chain length and their unsaturation, could have caused di(16:0)PC to be grossly over-represented relative to di(18:1)PC in vesicles made by our emulsion transfer method. Likewise, these characteristics could have contributed to the small shifts in PC-lipids in vesicles made by other techniques. To determine which characteristics cause major shifts in lipid ratios, we designed three sets of experiments using binary mixtures of lipids. The first set probes lipid unsaturation. The second probes lipid chain length. The third probes PC-lipids vs. sterol content. Vesicles made by gentle hydration required a small fraction of charged lipid (∼5 mol% di(18:1)PG).

The first set of binary experiments reveals that all methods except for emulsion transfer faithfully incorporate lipids from the stock solution (∼50%:50% di(16:0)PC/di(16:1)PC) into vesicles, within sample-to-sample variation. In contrast, saturated lipid is overrepresented in our emulsion technique, across two sets of experiments (Fig. 4, Fig. S15). This overrepresentation of saturated lipid by a factor of ∼1.2 cannot be due to known, favorable interactions between saturated lipids and cholesterol because the binary solution contains no cholesterol (51). Given that sample-to-sample lipid ratios in the 5-component vesicles in Fig. 3 were within ±2 mol%, we expect ratios in the binary mixtures (which have higher overall percentages) to be within ∼4 mol%, which generally holds true. Small sample sizes of *n* = 3 contribute to variations in uncertainties; uncertainties of ±1 mol% are observed in Fig. S15A, and uncertainties of ±5 mol% are observed in Fig. 4A, for the same lipid composition and method. Representative micrographs of vesicles appear in Fig. S12.

**Figure 4.**
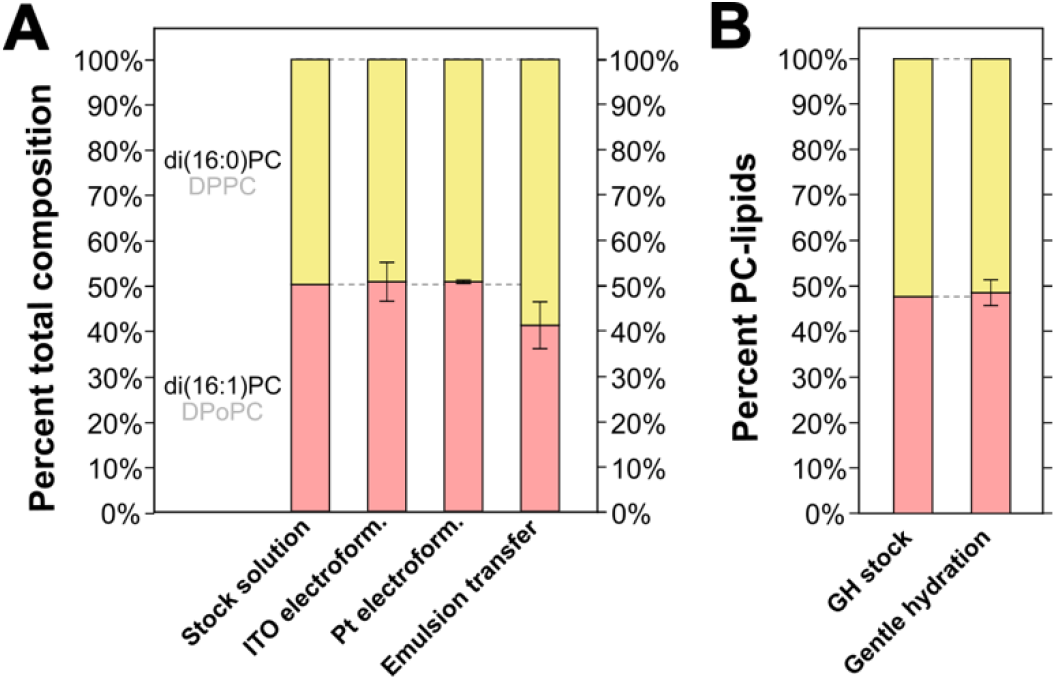
Percent of saturated and unsaturated lipids in binary vesicles produced by different methods. **A)** Percent of di(16:0)PC and di(16:1)PC in a stock solution and in averages from three independent vesicle solutions of electroformation and emulsion transfer techniques made from the stock. Error bars above each section of the bar chart are standard deviations of the three independent experiments. Full data appear in Tables S2, S3, and S13. **B)** Percent of di(16:0)PC and di(16:1)PC in the gentle hydration (GH) stock solution and averages from three independent gentle hydration vesicle solutions. This stock also contained charged lipids (≤5 mol% di(18:1)PG, see Fig. S16A).

The second set of binary experiments investigates the chain length of unsaturated PC-lipids (di(16:1)PC and di(18:1)PC; Fig. 5). Two methods (ITO electroformation and gentle hydration) faithfully incorporate the same ratio of these lipids from the stock solution. Pt wire electroformation incorporates slightly more long-chain lipid (a shift of ∼5 mol%). No conclusion can be reached from emulsion transfer experiments except that uncertainties are large, resulting in an apparent decrease of ∼2 mol% in long-chain lipid in one set of experiments (Fig. 5), counteracted by an increase of ∼6 mol% in an independent set of experiments (Fig. S15), with overlapping uncertainties. The large uncertainty is consistent with a lower yield of vesicles for this binary mixture by our emulsion transfer method, compared to other methods. Representative micrographs of vesicles appear in Fig. S13.

**Figure 5.**
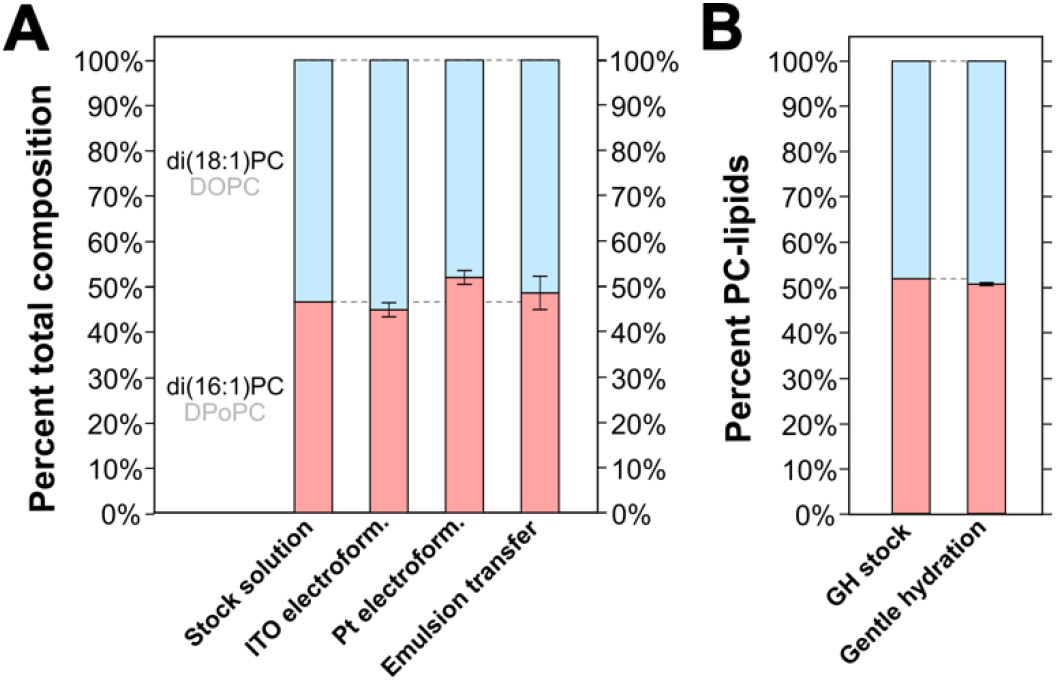
Percent of shorter and longer chained PC-lipids in binary vesicles produced by different methods. **A)** Percent of di(16:1)PC and di(18:1)PC in a stock solution and averages from three independent vesicle solutions of electroformation and emulsion transfer techniques made from the stock. Error bars above each section of the bar chart are standard deviations of the three independent experiments. Full data appear in Tables S4, S5, and S14. **B)** Percent of di(16:1)PC and di(18:1)PC in the gentle hydration (GH) stock solution and averages from three independent gentle hydration vesicle solutions. This stock also contained charged lipids (≤5 mol% di(18:1)PG, see Fig. S16B).

The third set of binary experiments (Fig. 6) investigates the mole fraction of cholesterol with respect to a PC-lipid (di(18:1)PC). Shifts in the lipid ratios mirror the trends in the 5-component vesicles in Fig. 3. For example, emulsion transfer again incorporates 80% less than the expected amount of cholesterol. Gentle hydration again incorporates 20% more than expected. This is surprising in light of recent results that find that cholesterol is, on average, slightly under-incorporated (about 10% less than expected) into vesicles of POPC (16:0/18:1PC) via a related gentle hydration technique (36). The remaining methods (ITO electroformation and Pt wire electroformation) again agree with the stock solution, within uncertainty. For all experiments, initial stock solutions were chosen to contain <50 mol% cholesterol, well below the maximum possible concentration of cholesterol in membranes (15, 18, 50). We chose to mix cholesterol with an unsaturated lipid because the mixture produced vesicles of high quality (Fig. S14), whereas the yield of vesicles comprised of cholesterol and saturated lipid (di(16:0)PC) were too low to accurately analyze.

**Figure 6.**
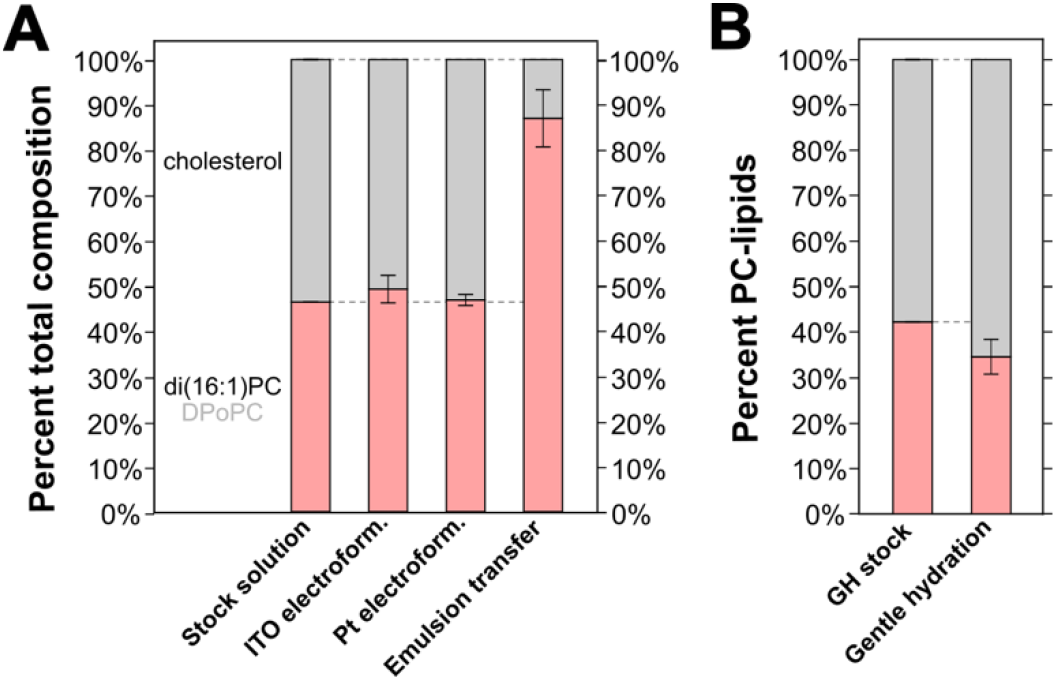
Percent of unsaturated PC-lipid and cholesterol in binary vesicles produced by different methods. **A)** Percent of di(16:1)PC and cholesterol in a stock solution and averages from three independent vesicle solutions of electroformation and emulsion transfer techniques made from the stock. Error bars above each section of the bar chart are standard deviations of the three independent experiments. Full data appear in Tables S6, S7, S15, and S16. **B)** Percent of di(16:1)PC and di(18:1)PC in the gentle hydration (GH) stock solution and averages from three independent gentle hydration vesicle solutions. This stock also contained charged lipids (≤5 mol% di(18:1)PG, see Fig. S16C).

**Figure 7.**
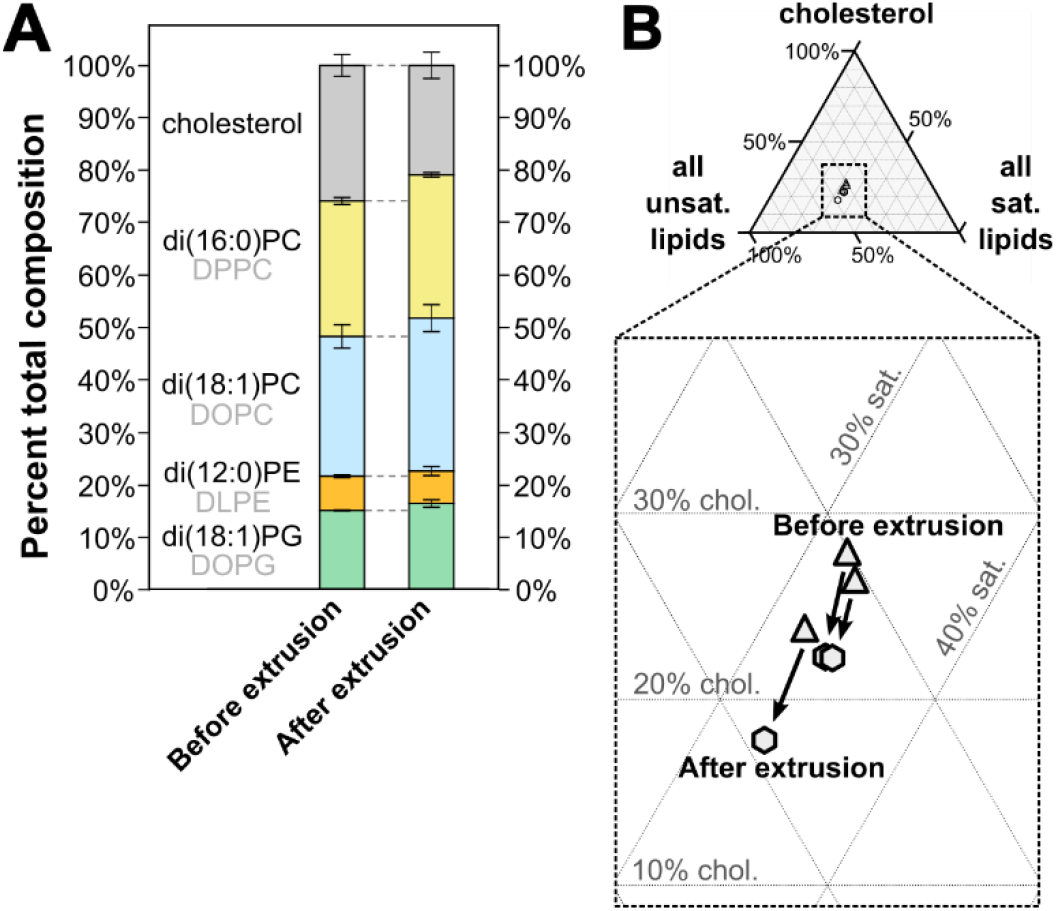
Extrusion of giant vesicles decreases their cholesterol content. **A)** Percentage of all five lipids in vesicles made via gentle hydration, before and after extrusion through a 0.1 µM filter. Values are averages, and error bars above each section of the bar chart are standard deviations of three independent experiments. **B)** Mole fractions of each experiment plotted on a pseudo-ternary diagram, before and after extrusion. The three vertices correspond to cholesterol, the sum of the saturated lipids (di(16:0)PC and di(12:0)PE), and the sum of the unsaturated lipids (di(18:1)PC and di(18:1)PG). Full data appear in Tables S8, S17, and S18. These gentle hydration samples were prepared and measured separately from samples comparing the four other methods.

### Extrusion of vesicles results in lower cholesterol fractions

Some powerful techniques of assessing membrane phase behavior use giant vesicles whereas others (e.g., electron microscopy and neutron scattering (52–54) require ∼100 nm vesicles, produced by extruding vesicles through polycarbonate filters. Here, we conducted a new series of experiments to compare ratios of lipids in solutions of giant vesicles generated by gentle hydration with ratios from the same solutions after extrusion. Corresponding micrographs are in Fig. S6.

We find systematic decreases in cholesterol due to extrusion (Fig. 5). In terms of absolute numbers, these shifts are not huge, a decrease of 5.0 ± 3.3 mol% cholesterol, where the uncertainty is the standard deviation for three independent experiments. In terms of the total percent of cholesterol, the shifts are significant, roughly 20% of cholesterol is lost upon extrusion of vesicles made by gentle hydration. This value falls in the middle of ranges previously reported: no loss of cholesterol was observed from some membranes (ternary mixtures of 2:1:3 PC-lipids/eggPC/cholesterol, binary mixtures of 33% cholesterol with di(18:1)PC or di(14:0)PC, or binary mixtures of 30/70, 40/60, or 50/50 cholesterol with 16:0/18:1PC), whereas a loss of ∼1/3 to ∼1/2 of cholesterol was observed from other membranes (for binary mixtures of 33% cholesterol with 16:0/18:1PC, di(16:0)PC, or di(20:4)PC (19, 36). In other words, no clear correlation seems to exist between lipid unsaturation and a loss of cholesterol from vesicles upon extrusion.

We find that the decrease in cholesterol upon extrusion is offset by smaller increases in PC-lipids (1.4 ± 0.7 mol% for di(16:0)PC, and 1.5 ± 0.8 mol% for di(18:1)PC, respectively). Shifts in PE-and PG-lipids are insignificant (0.4 ± 0.9 mol% for di(12:0)PE and 2.5 ± 3.4 mol% for di(18:1)PG). As before, one main conclusion is that experiments that report results at high precision (e.g., 1 mole%) will be sensitive to how vesicles are made and to sample-to-sample differences.

### Caveats

In the sections above, we tested whether several methods of making vesicles skew the ratio of lipids in membranes. Some variations on these methods have already been explored. For example, it was already known that varying the type of oil in emulsion transfer methods results in wide ranges of cholesterol content in vesicles (20–22, 55). Similarly, it was known that accurate membrane phase transitions (and presumably lipid compositions) relied on electroformation being performed at a temperature well above the highest melting temperature of any lipid in the sample using lipid films of sufficient thickness (23, 56).

However, many other variations on each method remain unexplored. For example, gentle hydration could have been conducted in salt solutions (rather than in sugar), or from lipid films deposited on surfaces of tracing paper (57) or polymer-coated substrates (58, 59) (rather than on bare glass). Similarly, gentle hydration of lipid films could have been achieved for different lengths of time using lipids in different ratios (36), followed by either shaking or vortexing (1) (rather than in a static solution) using different types of vortexers (36), and the subsequent multilamellar vesicles could have been made smaller through freeze/thaw cycles or sonication (rather than extrusion) (36, 52, 60). Details of each technique vary from laboratory to laboratory, and it is unclear how these details influence the ratio of lipids in membranes.

Similar caveats apply to the choice of stock solutions. Many lipid mixtures are relevant, and each may produce different shifts in lipid ratios, especially if interdependencies between lipids affect their incorporation into vesicles. These interdependencies will be challenging to assess when lipid stocks contain small fractions of some lipids (like the PE-and PG-lipids in our 5-component sample) because correspondingly small shifts in their absolute mole fractions will be observed, even if percent changes are high. Although we found minor shifts in lipid fractions for most methods, it is worth bearing in mind that small shifts can result in big changes in membrane behavior. For example, if a membrane lies near a phase boundary, a small shift in its lipid composition can lead a previously uniform membrane to demix into coexisting phases, or a liquid membrane to turn into a gel phase. For researchers who are comparing vesicles made by different methods, a best practice is to choose a ratio of lipids that lies in the middle of the preferred phase region.

## CONCLUSIONS

In summary, the five methods we used to make vesicles result in a range of deviations in lipid ratio relative to 5-component and binary stock solutions. Electroformation, whether on an ITO slide or on platinum wires, results in the smallest shifts (≤5 mole%). Researchers will find the magnitudes of these shifts to be either comfortingly small or alarmingly large, depending on how they design their experiments. For example, experiments that report concentrations incremented by only ∼1 mole% will be highly sensitive to the choice of vesicle-making method and to sample-to-sample differences. On the other side of the spectrum, the largest shifts are observed with emulsion transfer. It was already known that emulsion transfer resulted in a wide range of low cholesterol fractions in vesicles (20–22, 55). In our experiments, ∼80% less cholesterol was incorporated into vesicles than expected from the stock solution. A surprising result is that emulsion transfer also shifts the relative amount of saturated and unsaturated lipid; it incorporates more di(16:0)PC than expected relative to di(16:1)PC.

For the 5-component stock, sample-to-sample variations for all methods were roughly ±2 mol% for independent experiments. Again, researchers who rely on 1% precision will be sensitive to sample-to-sample variations, and others will find the reproducibility reassuring. As mole fractions increase for each lipid, some corresponding sample-to-sample variations increase concomitantly (as in binary vesicles made by emulsion transfer), whereas others remained low, on the order of ±2 mol%.

The shifts in lipid compositions that we measured are context dependent. That context can include the types of lipids in the system (as explored in the binary mixtures in Figs. 4-6 and S15), the amount of lipid, the type of solvents, the temperature, and differences in experimental protocol. What should communities of researchers do when faced with so many potential variables? Of course, we should adhere to accepted best practices (e.g., electroforming with enough lipid and at high enough temperature) and we should report detailed descriptions of our methods. We can shift our mindset to think big rather than to think small. Armed with estimates of how lipid compositions might vary from lab to lab, we have a better sense of how to look for broad trends and new phenomena, and to celebrate when those trends are supported by other methods, even if the quantitative data vary subtly.

## AUTHOR CONTRIBUTIONS

All authors designed research and protocols. HMJW, KJW, GJG, ELP, and AL performed research and analyzed data. SLK, HMJW, KJW, and GJG wrote the manuscript. All authors edited the manuscript.

## ACKNOWLEDGEMENTS

We thank the staff at the UW Statistical Consulting Services for their expertise. HMJW was supported by the National Institute of General Medical Sciences of the National Institutes of Health under Award Number T32GM008268. This research was supported by National Science Foundation grant MCB-1925731 and MCB-2325819 to SLK and by National Institutes of Health grant R01HD092659 and R01 AI136979 to LX.

## Supporting Material for

**Figure S1.**
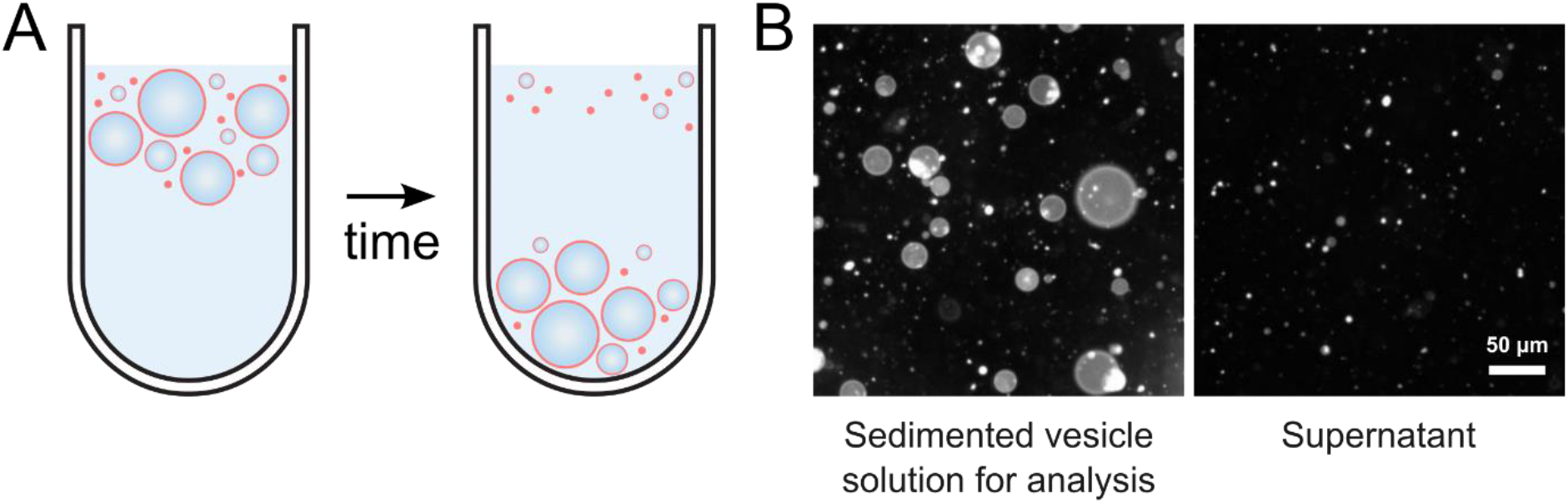
Sedimenting vesicles removes some lipid aggregates. **A)** Vesicles filled with a dense sucrose solution sink in an osmotically matched glucose solution. **B)** After sinking, large vesicles are observed only in the sedimented solution, and not in the supernatant. The vesicles in these images were made by emulsion phase transfer. The low density of giant vesicles in the supernatant is representative of all techniques.

**Figure S2.**
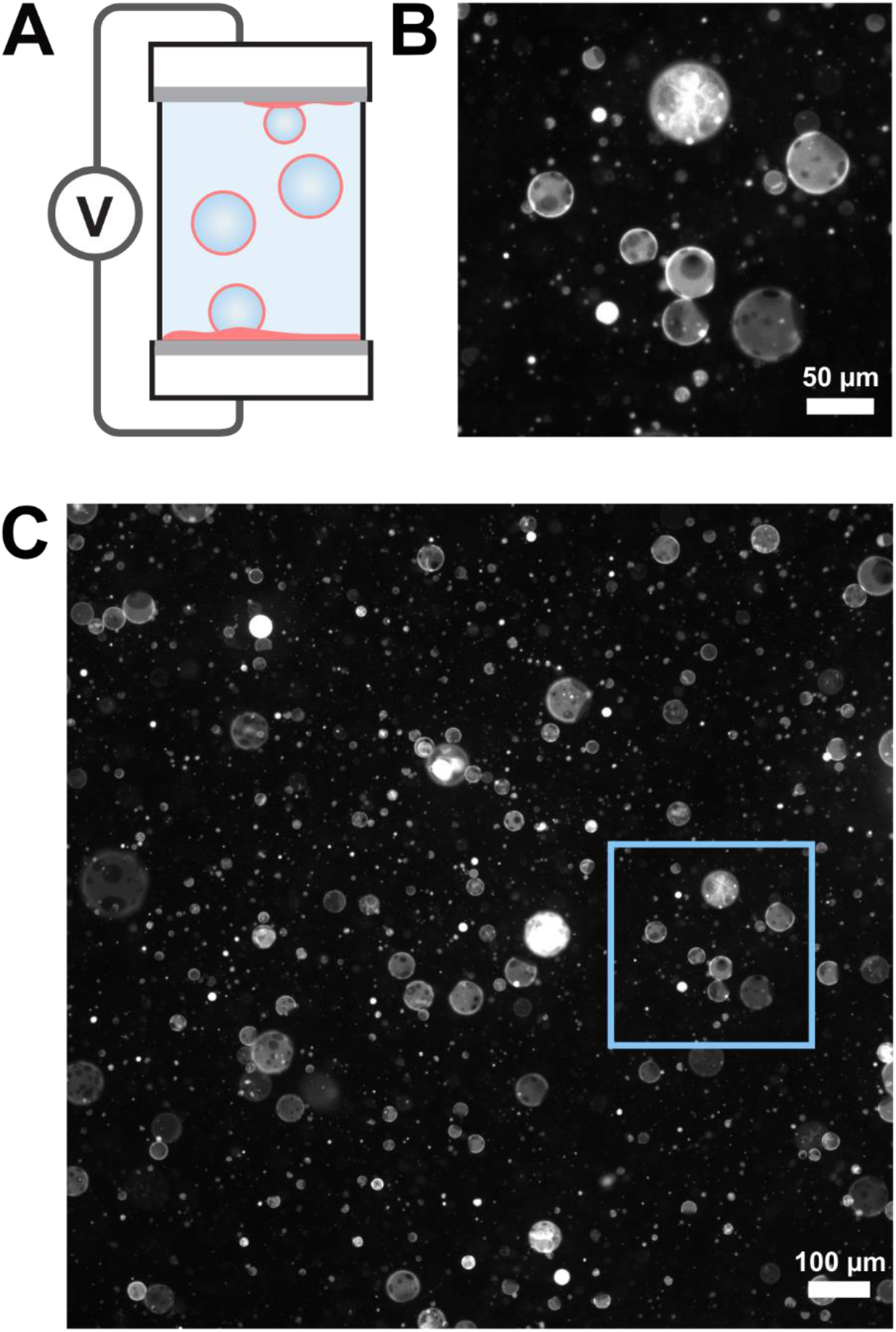
Representative fluorescence micrographs of vesicles made by electroformation on ITO-coated slides. **A)** Schematic of ITO electroformation. **B)** Close-up of vesicles in the blue box in panel C. **C)** Larger field of view of vesicles produced by electroformation on ITO slides. As expected, ITO electroformation produces the highest yield of giant, unilamellar vesicles (GUVs) of the four methods tested for producing these vesicles. Some multilamellar vesicles, some nested vesicles (large vesicles filled with smaller vesicles), and some bright, lipid aggregates are also produced.

**Figure S3.**
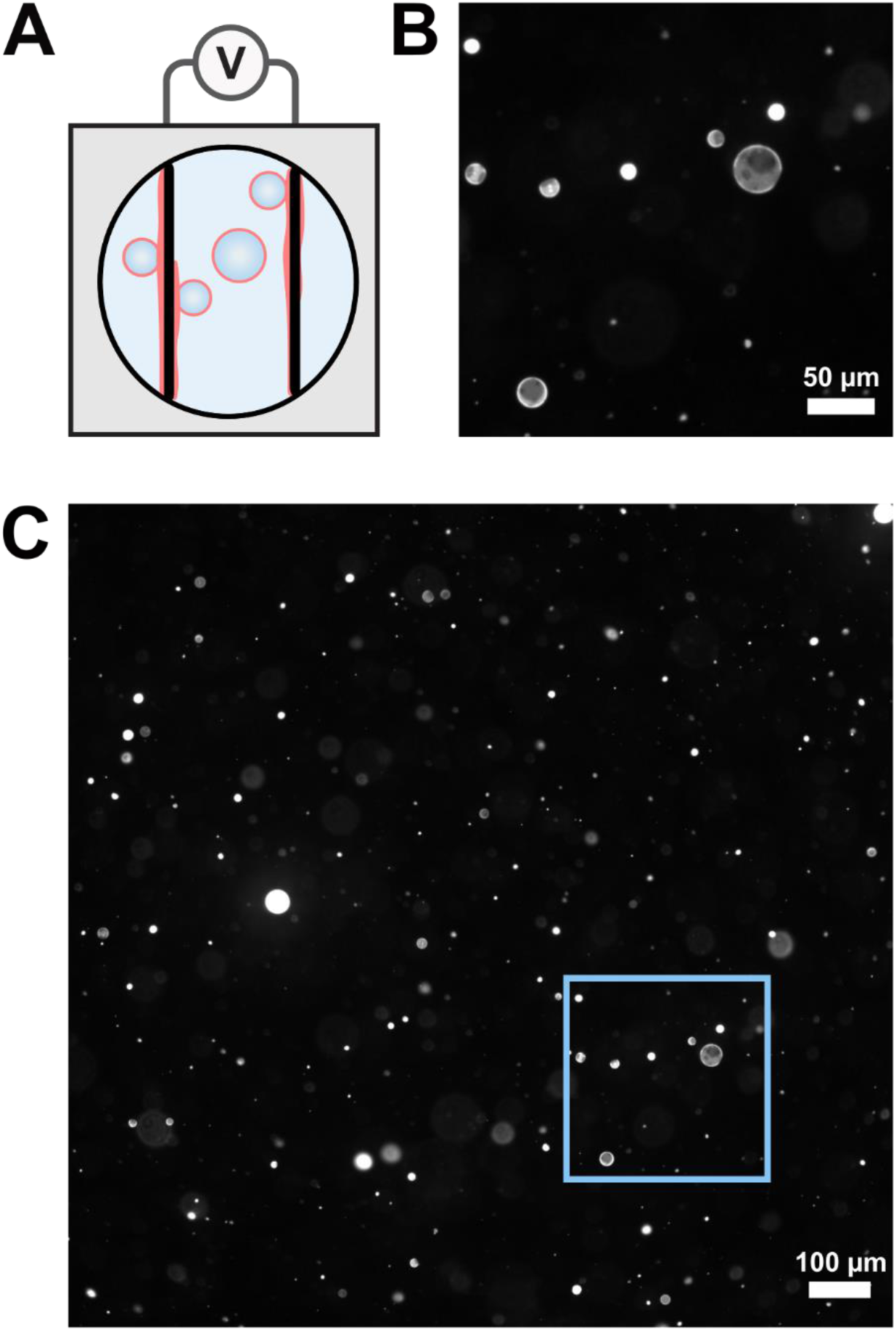
Representative fluorescence micrographs of vesicles made by electroformation on platinum wires. **A)** Schematic of Pt electroformation. **B)** Close-up of vesicles in the blue box in panel C. **C)** Larger field of view of vesicles produced by electroformation on Pt wires. The volume of stock solution used is low relative to the other methods and therefore produces a lower yield of vesicles at the same stock concentration.

**Figure S4.**
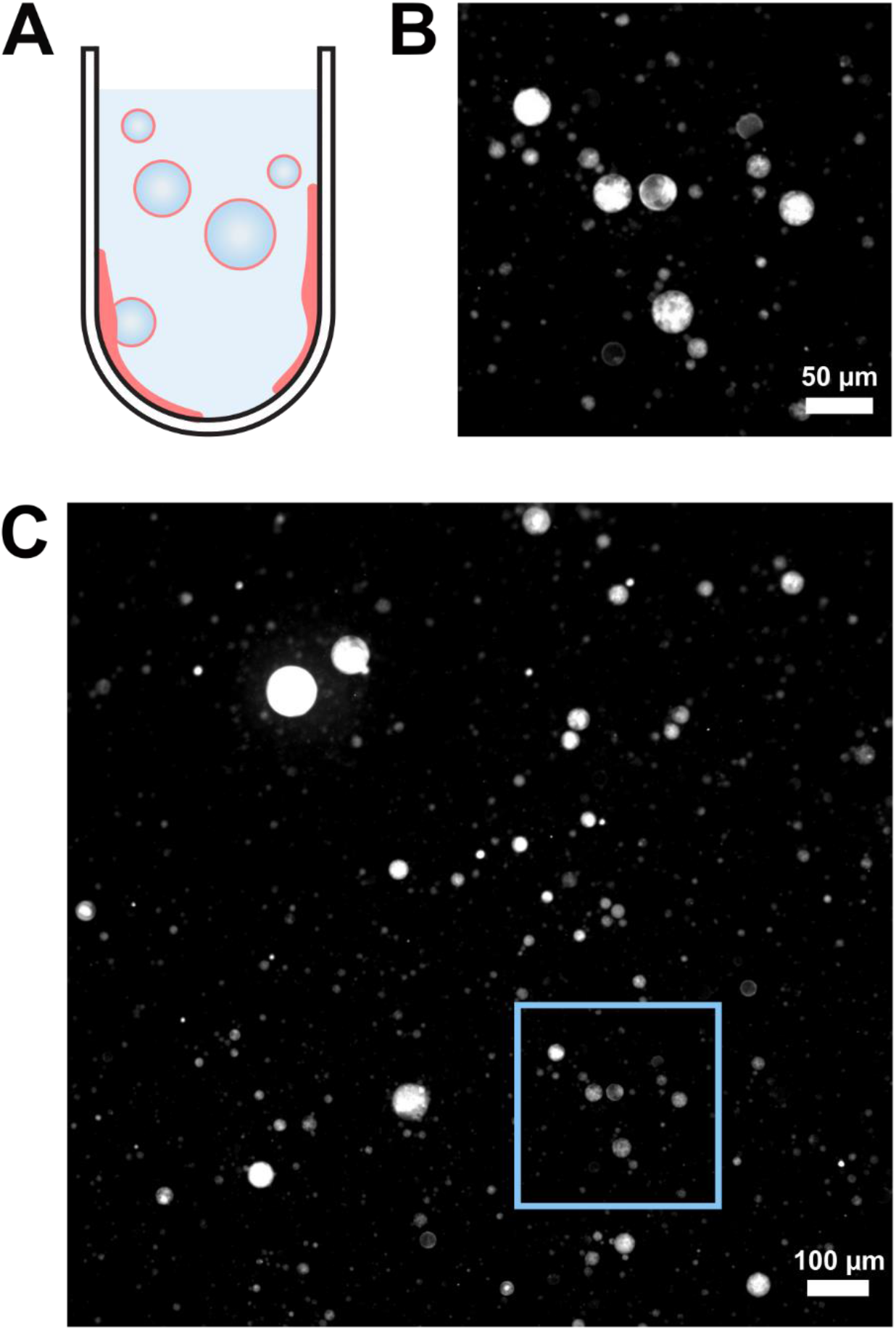
Representative fluorescence micrographs of vesicles made by gentle hydration for 24 hrs. **A)** Schematic of gentle hydration. **B)** Close-up of vesicles in the blue box in panel C. **C)** Larger field of view of vesicles produced by gentle hydration. As expected, the sample predominantly contains multilamellar vesicles and nested vesicles. When contrast is optimized for the bright, multi-layered vesicles, unilamellar vesicles in the same field of view can be too dim to be imaged.

**Figure S5.**
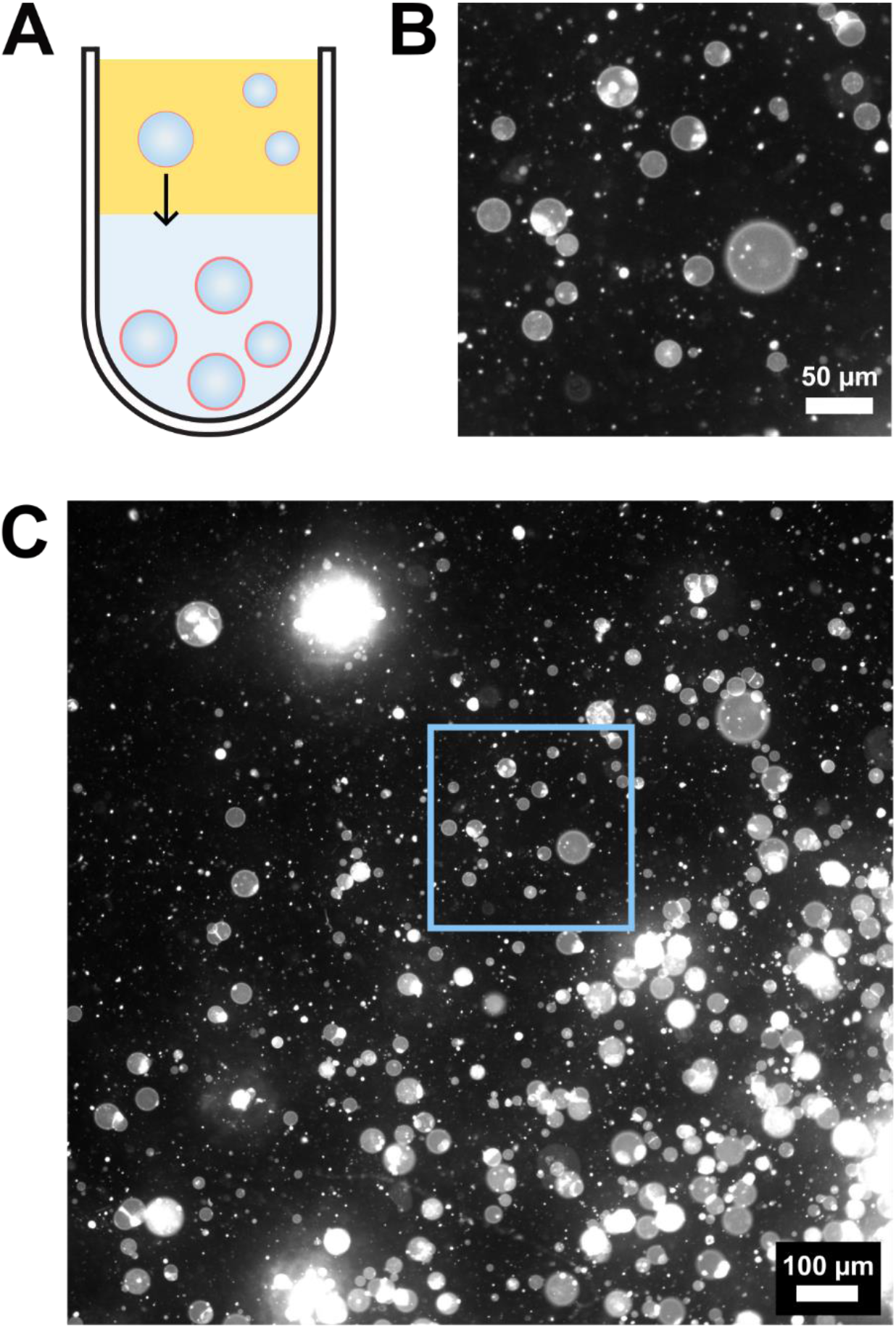
Representative fluorescence micrographs of vesicles made by emulsion phase transfer. **A)** Schematic of emulsion phase transfer. **B)** Close-up of vesicles in the blue box in panel C. **C)** Larger field of view of vesicles produced by emulsion phase transfer. The emulsion transfer method produces a high yield of giant unilamellar vesicles as well as defects including nested vesicles and lipid aggregates.

**Figure S6.**
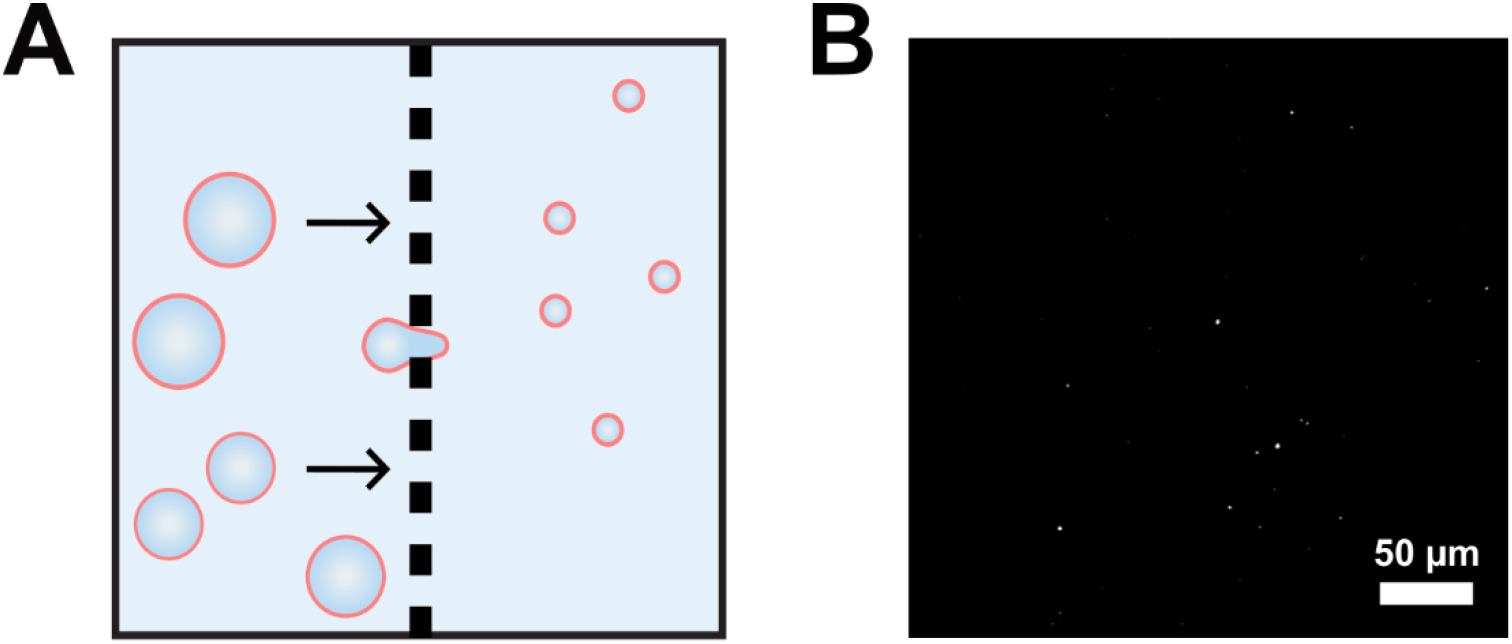
Vesicles after extrusion. **A)** Schematic of extrusion. **B)** Representative fluorescence micrograph of vesicles after extrusion through a filter with 100 nm diameter holes. As expected, most extruded vesicles are too small to be resolved with standard fluorescence microscopy techniques.

**Figure S7.**
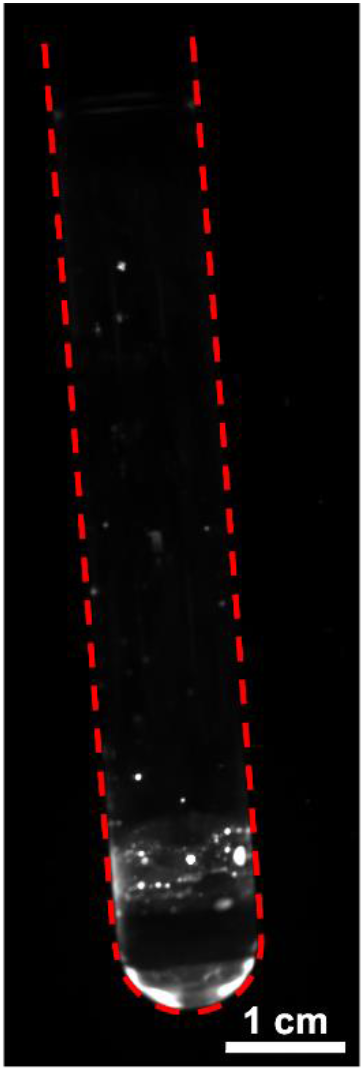
Residual lipid due to the gentle hydration technique, detected by fluorescence microscopy. Residual lipid left on the interior of a glass test tube is shown after a sample of vesicles made by gentle hydration was removed. A dashed red outline defines the edges of the test tube. The image was captured on an Amersham ImageQuant 800 (Cytiva) with a Cy3 (UV) filter.

**Figure S8.**
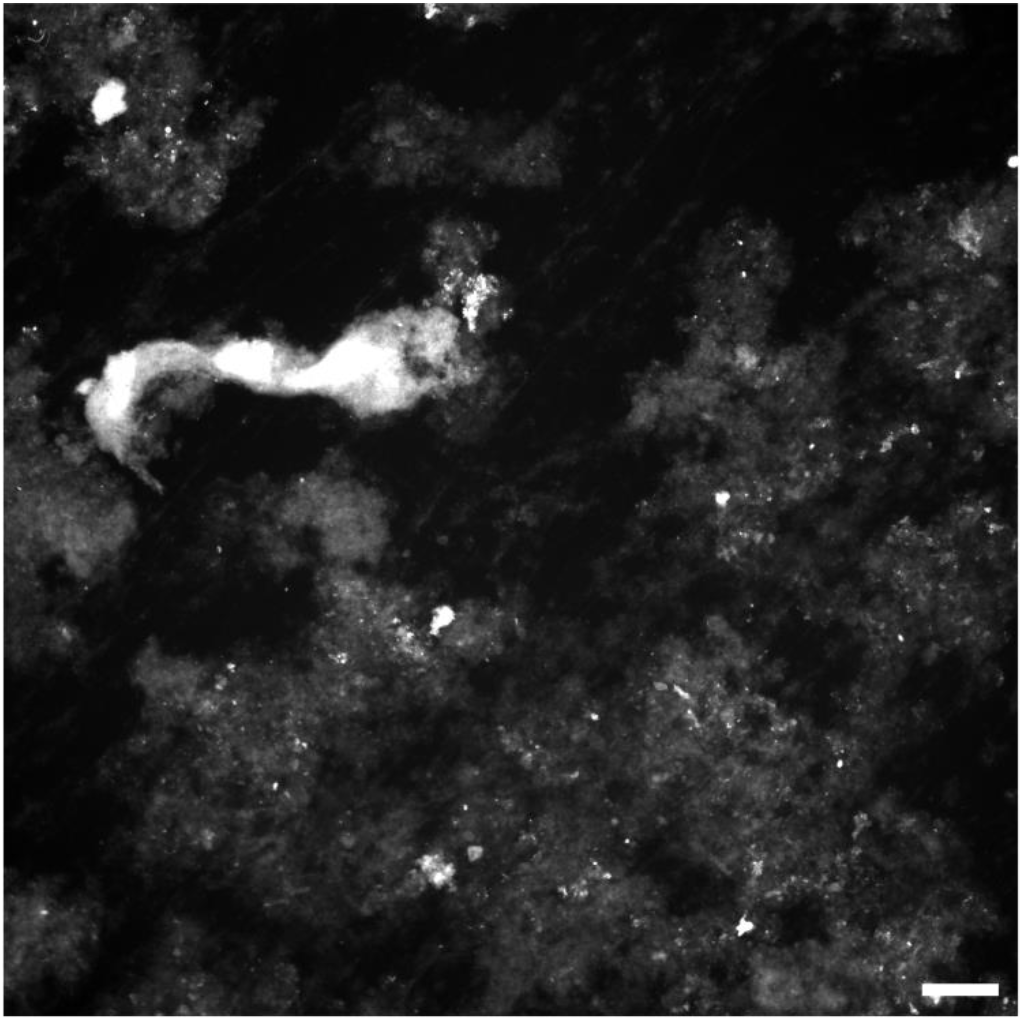
Residual lipid due to the ITO electroformation technique, detected by fluorescence microscopy. Residual lipid left on an ITO slide is shown after a sample of vesicles made by electroformation was removed. The image is representative of all experiments using electroformation on ITO slides. Scale bar: 500 μm.

**Figure S9.**
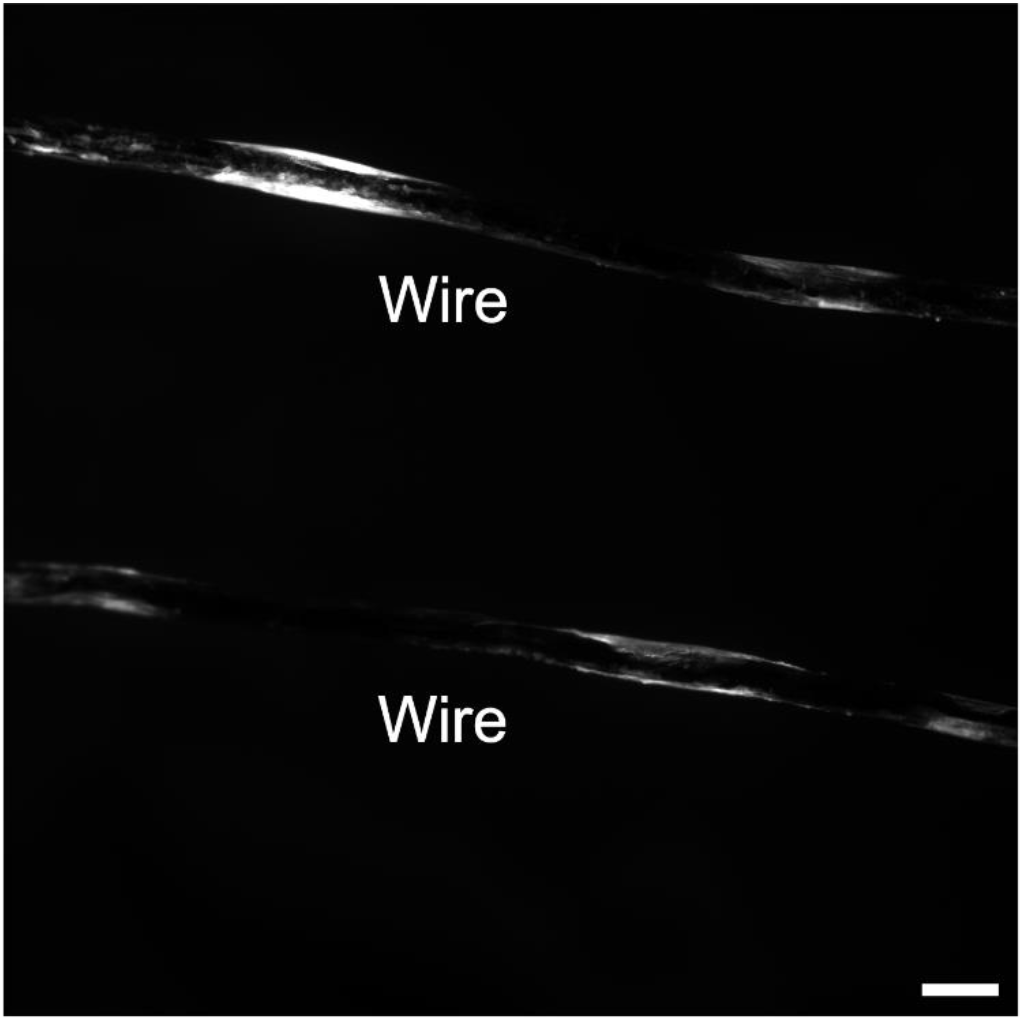
Residual lipid due to the Pt wire electroformation technique, detected by fluorescence microscopy. Residual lipid left on platinum wires is shown after a sample of vesicles made by electroformation was removed. The image is representative of all experiments using electroformation on Pt wires. Scale bar: 500μm.

**Figure S10.**
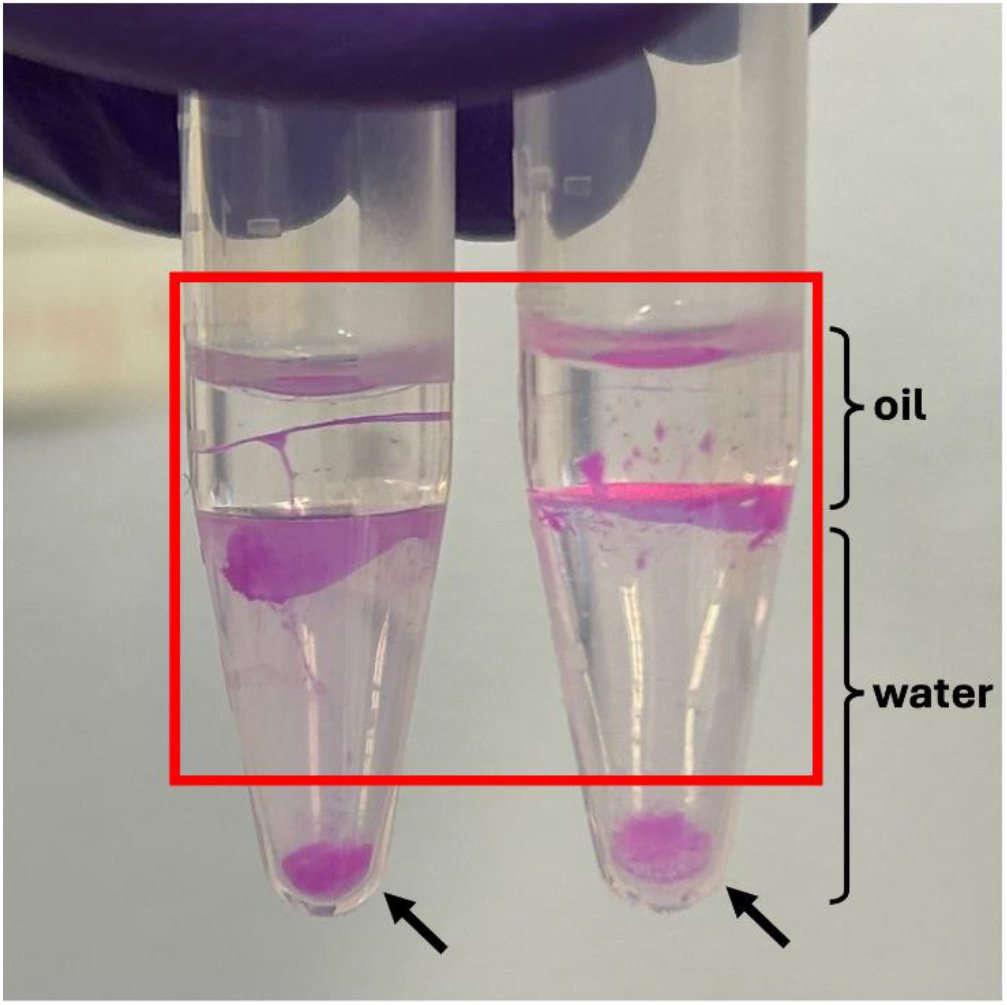
Residual lipid due to the emulsion transfer method, imaged in ambient light. Residual lipid left in oil and at the oil-water interface is shown after completing two emulsion transfer experiments. The images are representative of all experiments using emulsion transfer. The red box encloses the supernatant, which was removed before analysis by mass spectrometry. Most residual lipid is trapped at the interface between the water and oil. Black arrows indicate pellets of vesicles. These pellets were subsequently resuspended in 300 mM glucose solution, imaged, and analyzed as described in the methods.

**Figure S11.**
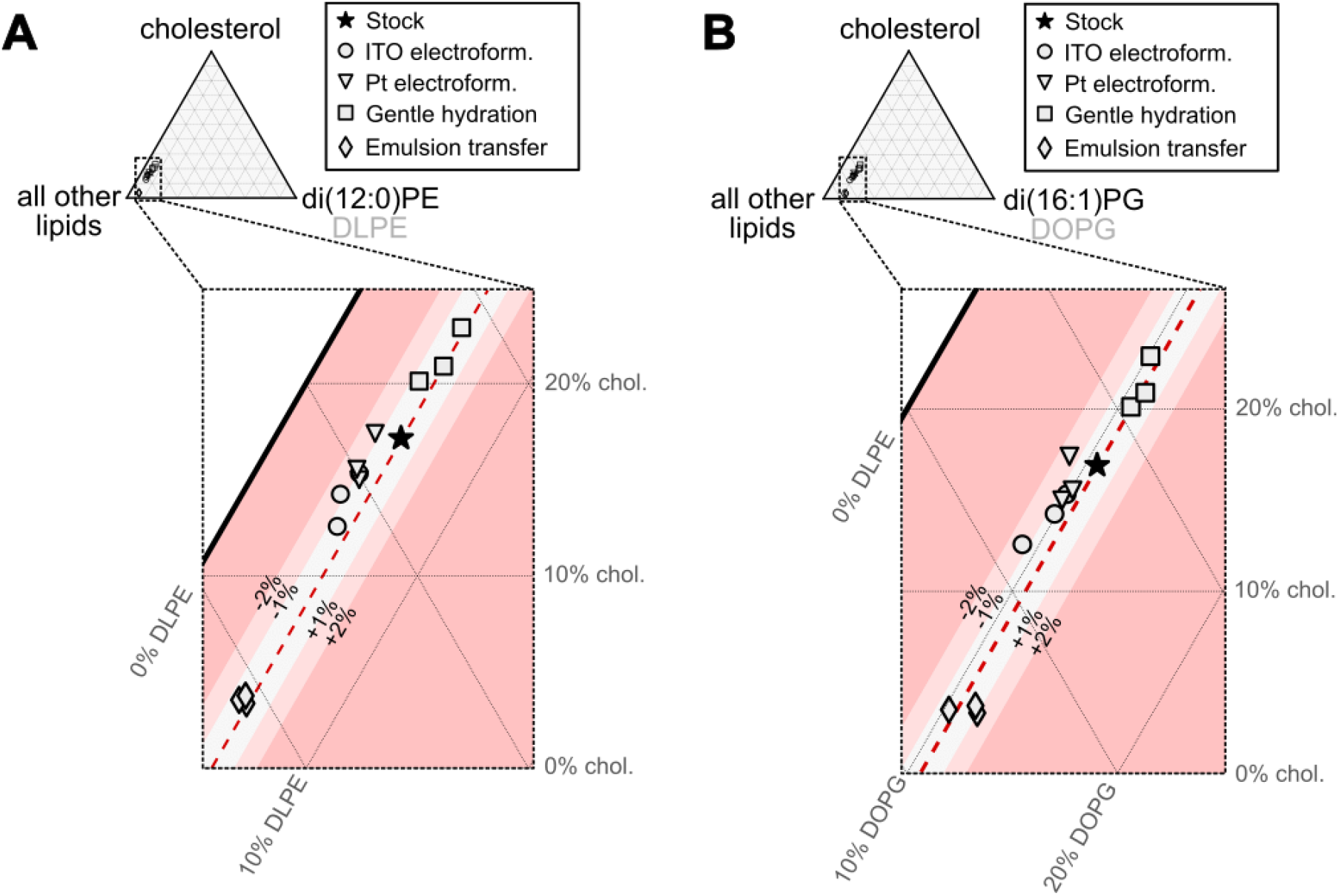
Lipid percentages in 5-component vesicles produced by different methods, plotted as pseudo-ternary diagrams in which one vertex is di(12:0)PE (DLPE) or di(18:1)PG (DOPG). Vesicles were prepared by four methods: electroformation on ITO slides, electroformation on platinum wires, gentle hydration, and emulsion phase transfer. Three independent preparations and experiments were run for each method. **(A)** Lipid percentages from each independent experiment are plotted on a pseudo-ternary diagram, where the three vertices represent di(12:0)PE, cholesterol, and the sum of all other lipids. **(B)** Lipid percentages from each independent experiment are plotted on a pseudo-ternary diagram, where the three vertices represent di(18:1)PE, cholesterol, and the sum of all other lipids. In both panels, the lipid composition measured for the stock solution is shown as a star. Deviations from the master stock solution in increments of 1% and 2% of DLPE or DOPG are shown by shaded bands.

**Figure S12.**
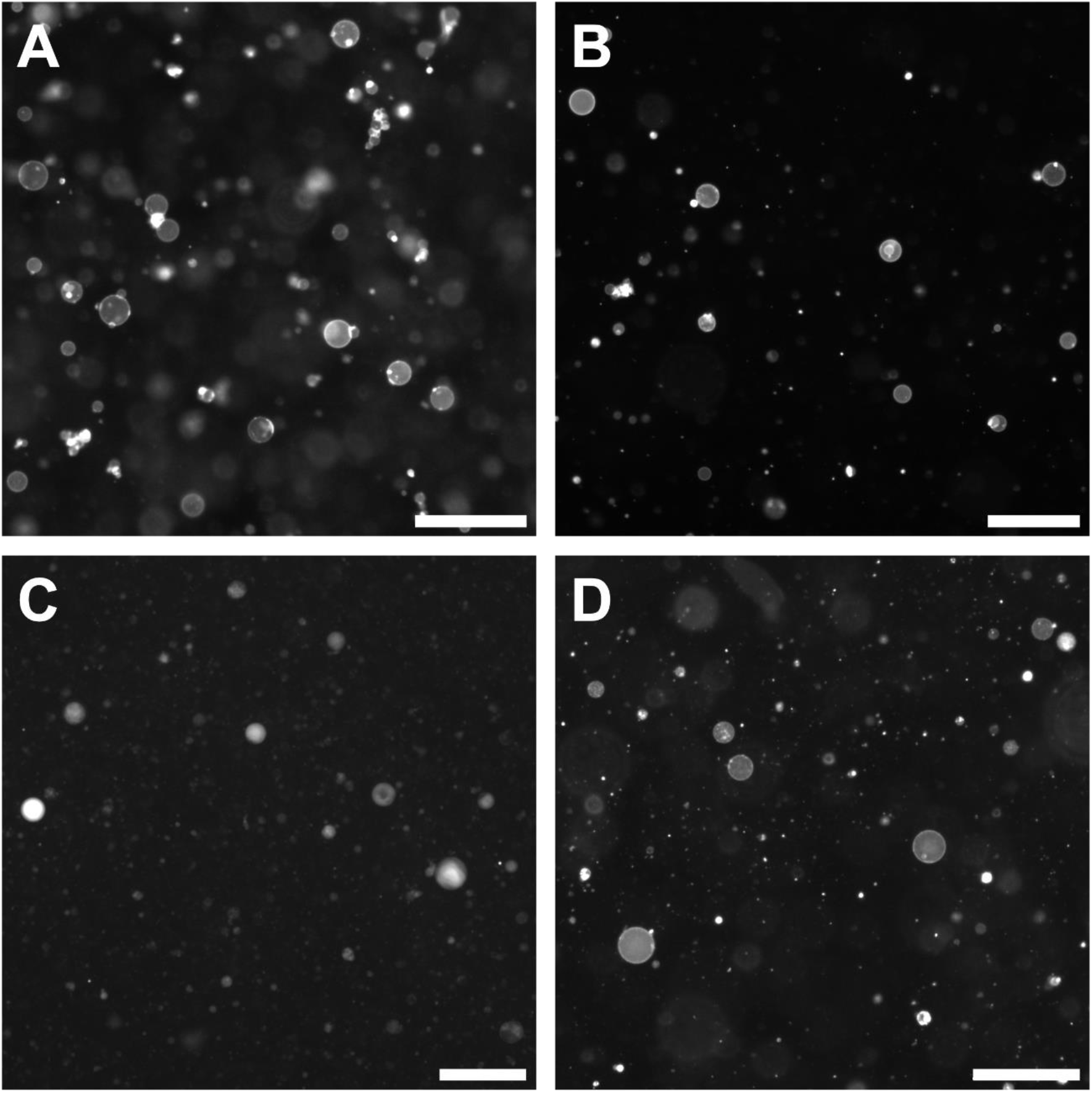
Representative fluorescence micrographs of solutions of vesicles comprised of binary mixtures of saturated lipid (di(16:0)PC) and unsaturated lipid (di(16:1)PC). Vesicles were formed by **A)** ITO electroformation, **B)** Pt wire electroformation, **C)** gentle hydration, and **D)** emulsion transfer. For this lipid composition, most giant unilamellar vesicles have some type of defect such as adhered vesicles, nested vesicles, or bright aggregates (Panels A, B, and D). Scale bars: 100 μm.

**Figure S13.**
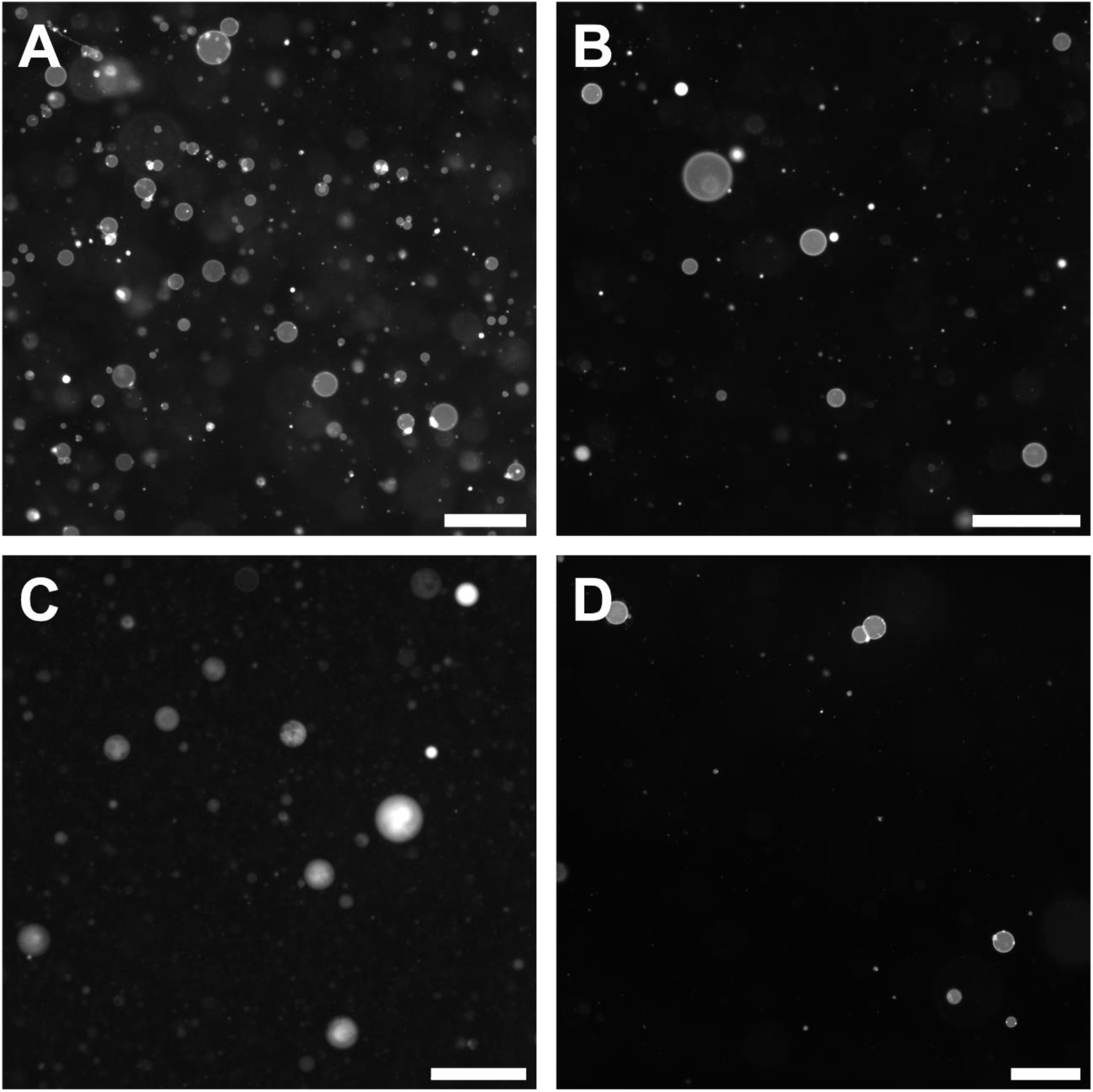
Representative fluorescence micrographs of solutions of vesicles comprised of binary mixtures of unsaturated lipids with shorter chains (di(16:1)PC and longer chains (di(18:1)PC). Vesicles were formed by **A)** ITO electroformation, **B)** Pt wire electroformation, **C)** gentle hydration, and **D)** emulsion transfer. For this mixture of lipids, emulsion transfer had lower vesicle yields compared to other techniques. Scale bars: 100 μm.

**Figure S14.**
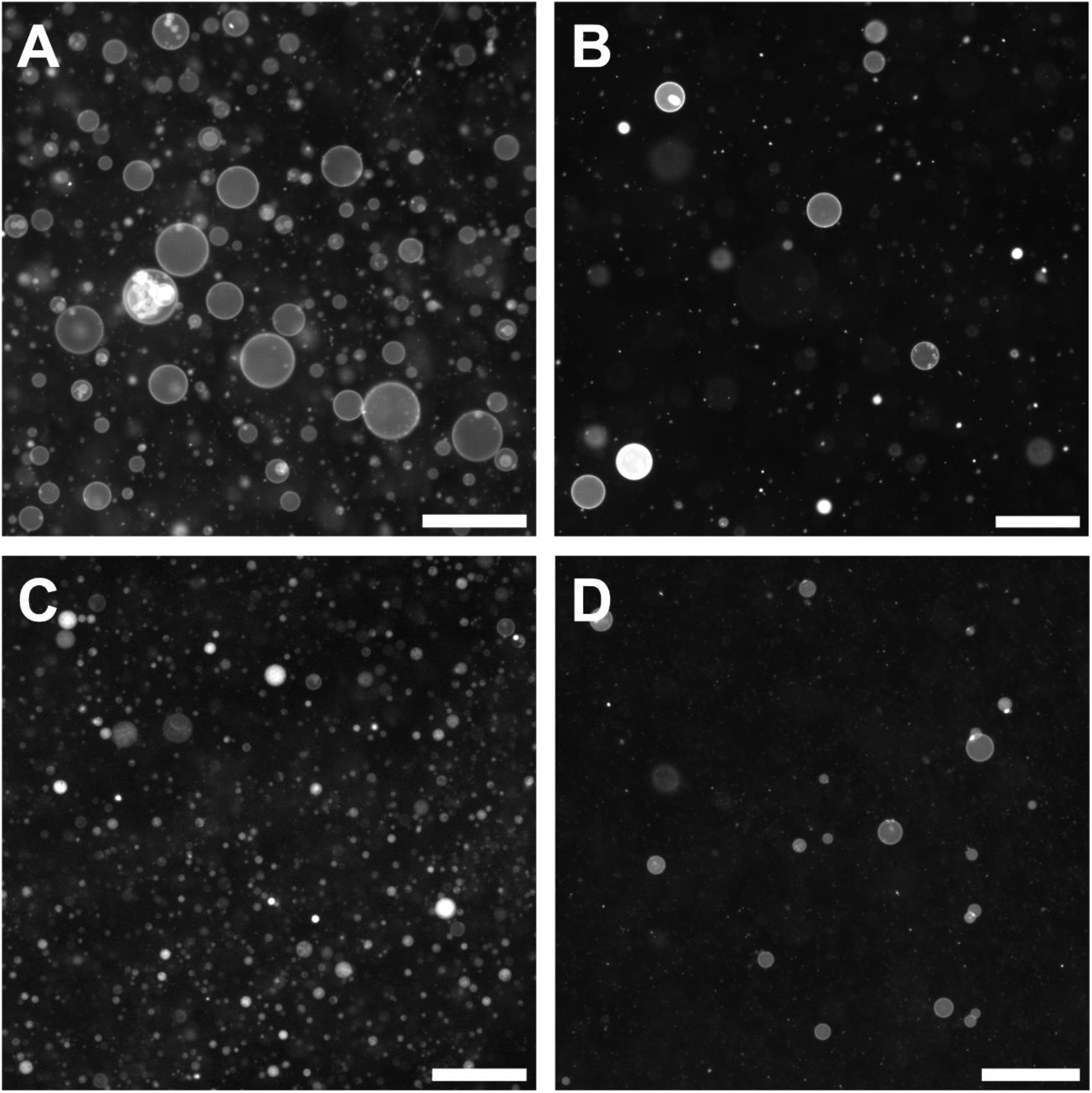
Representative fluorescence micrographs of solutions of vesicles comprised of binary mixtures of an unsaturated lipid (di(16:1)PC) and cholesterol. Vesicles were formed by **A)** ITO electroformation, **B)** Pt wire electroformation, **C)** gentle hydration, and **D)** emulsion transfer. For this mixture of lipids, gentle hydration yielded smaller vesicles compared to other techniques. Scale bars: 100 μm.

**Figure S15.**
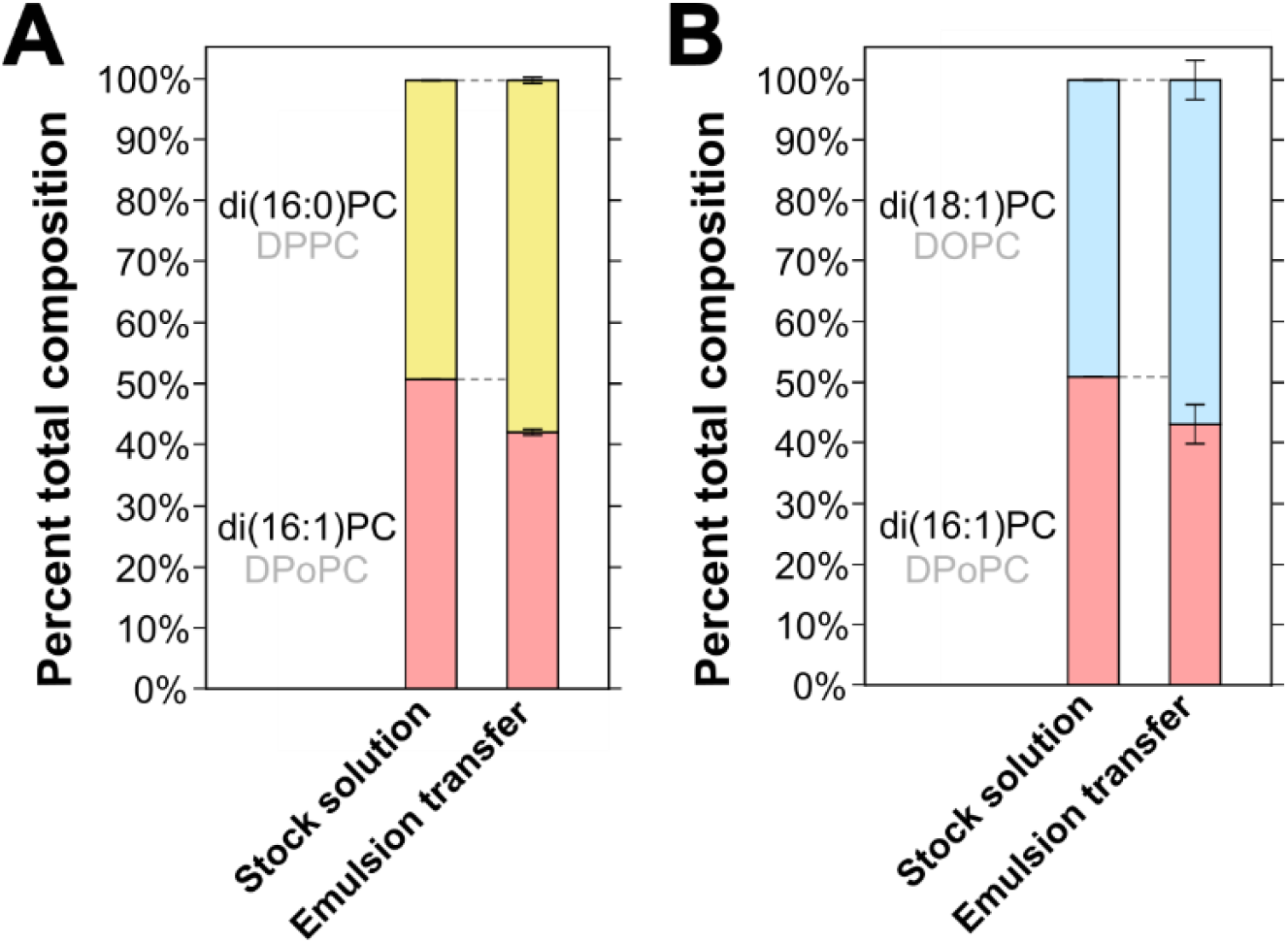
Saturated lipids and longer-chain lipids are over-represented in binary vesicles made by emulsion transfer. **A)** Percent of a saturated PC-lipid (di(16:0)PC) and an unsaturated lipid (di(16:1)PC) in a stock solution and in vesicles made by emulsion transfer from that stock. **B)** Percent of a longer-chain PC-lipid (di(18:1)PC) and a shorter-chain lipid (di(16:1)PC) in a stock solution and in vesicles made by emulsion transfer from that stock. In both panels, values are averages of three independent experiments, error bars above each section of the bar chart are standard deviations, and all experiments are independent from the data in Figs. 4 and 5 of the main text, including the use of a different stock solution. For these triplicates, it was not necessary to concentrate them by a factor of 5 since abundances were sufficiently above the background. Full data appear in Tables S9, S10, S19 and S20.

**Figure S16.**
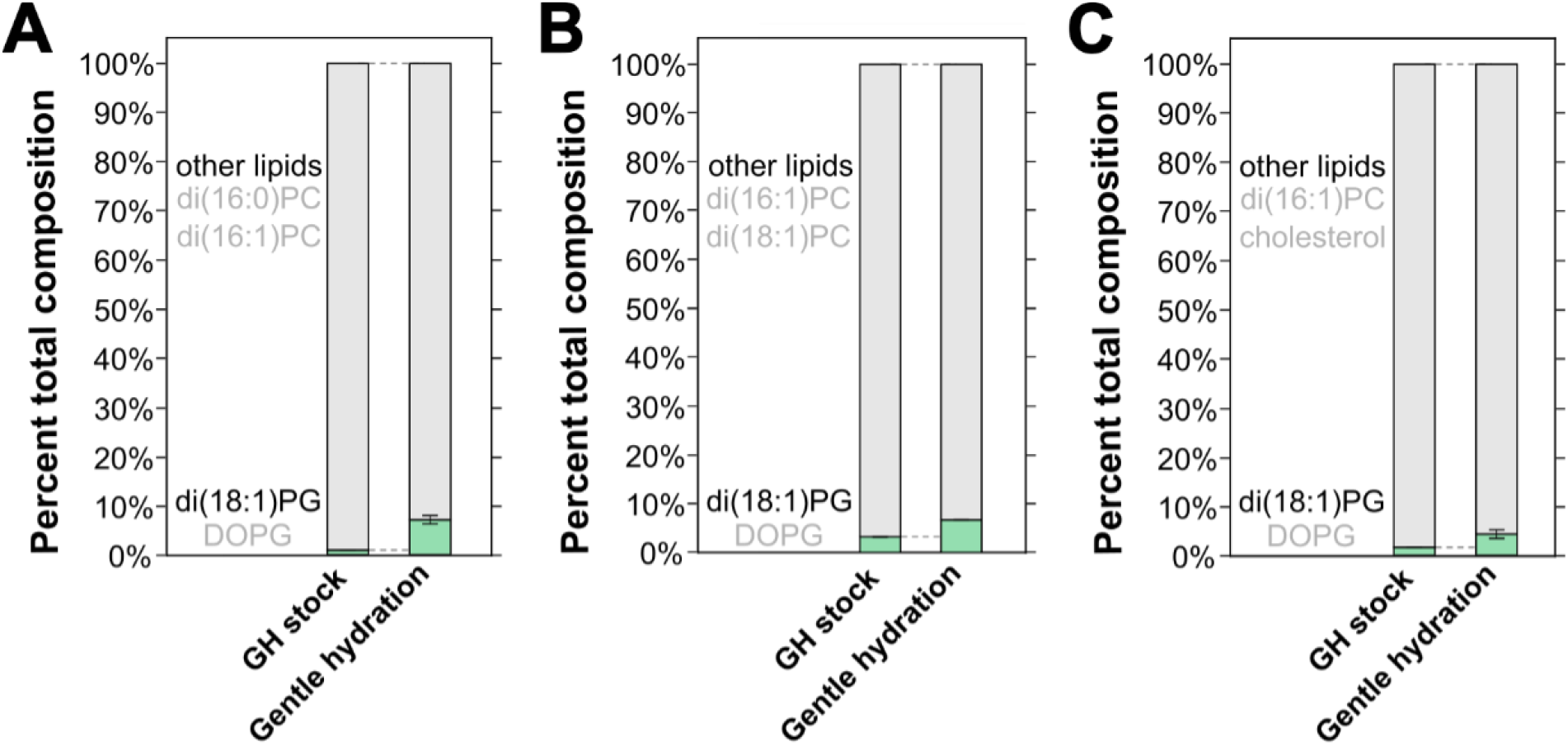
PG-lipids are over-represented in vesicles made by gentle hydration from stock solutions initially containing <5 mol% PG-lipid. In all three panels, the percent of di(18:1)PC lipids and the percent of other lipids are shown for a stock solution in chloroform (left bar of each panel) and for vesicles made by gentle hydration from those stocks (right bar). Vesicle data are averages, and error bars above each section of the bar chart are standard deviations of three independent experiments. **A)** The other lipids are di(16:0)PC and di(16:1)PC. Full data appear in tables S3 and S13. **B)** The other lipids are di(16:1)PC and di(18:1)PC. Full data appear in tables S5 and S14. **C)** The other lipids are di(16:1)PC and cholesterol. Full data appear in tables S7, S15, and S16.

**Figure S17.**
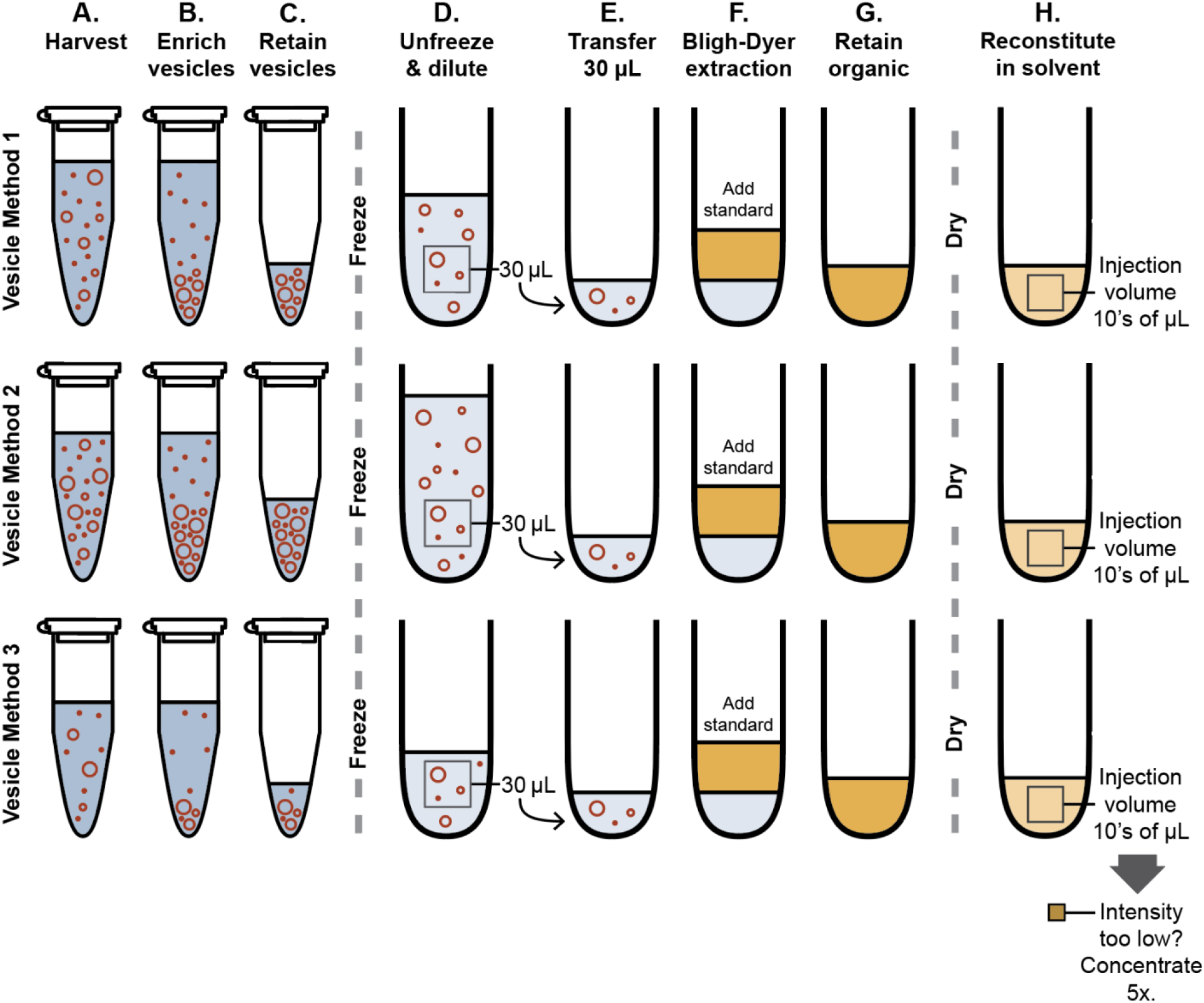
Extraction Procedure. **A)** Each vesicle method produces an aqueous solution containing vesicles and lipid aggregates. Concentrations and volumes of each solution may be different. **B)** A portion of each solution is enriched in sedimented vesicles. **C)** The portion of each solution enriched in vesicles is retained and frozen. The remainder is discarded. **D)** Batches of samples are unfrozen at the same time and diluted. The volume for dilution is calculated with an assumption that 10% of the stock lipids used to make vesicles in each experiment is retained in Step C and with a goal that the concentration of the least prevalent lipid is ∼1 µM in Step G. **E)** An aliquot of 30 µL from each sample is transferred to new glassware. **F)** Lipid standards are added to each sample, and all lipids are extracted into an organic solvent. **G)** The organic phase is retained and the aqueous phase is discarded. **F)** The lipids are dried and then reconstituted in 2:1 acetonitrile:methanol. Tens of microliters of this solution is injected into the HILIC-IM-MS apparatus. If the intensity the peak of the least prevalent lipid is too low, the remaining solution is concentrated 5 times and the same volume of solution is injected into the HILIC-IM-MS apparatus.

**Table S1.**
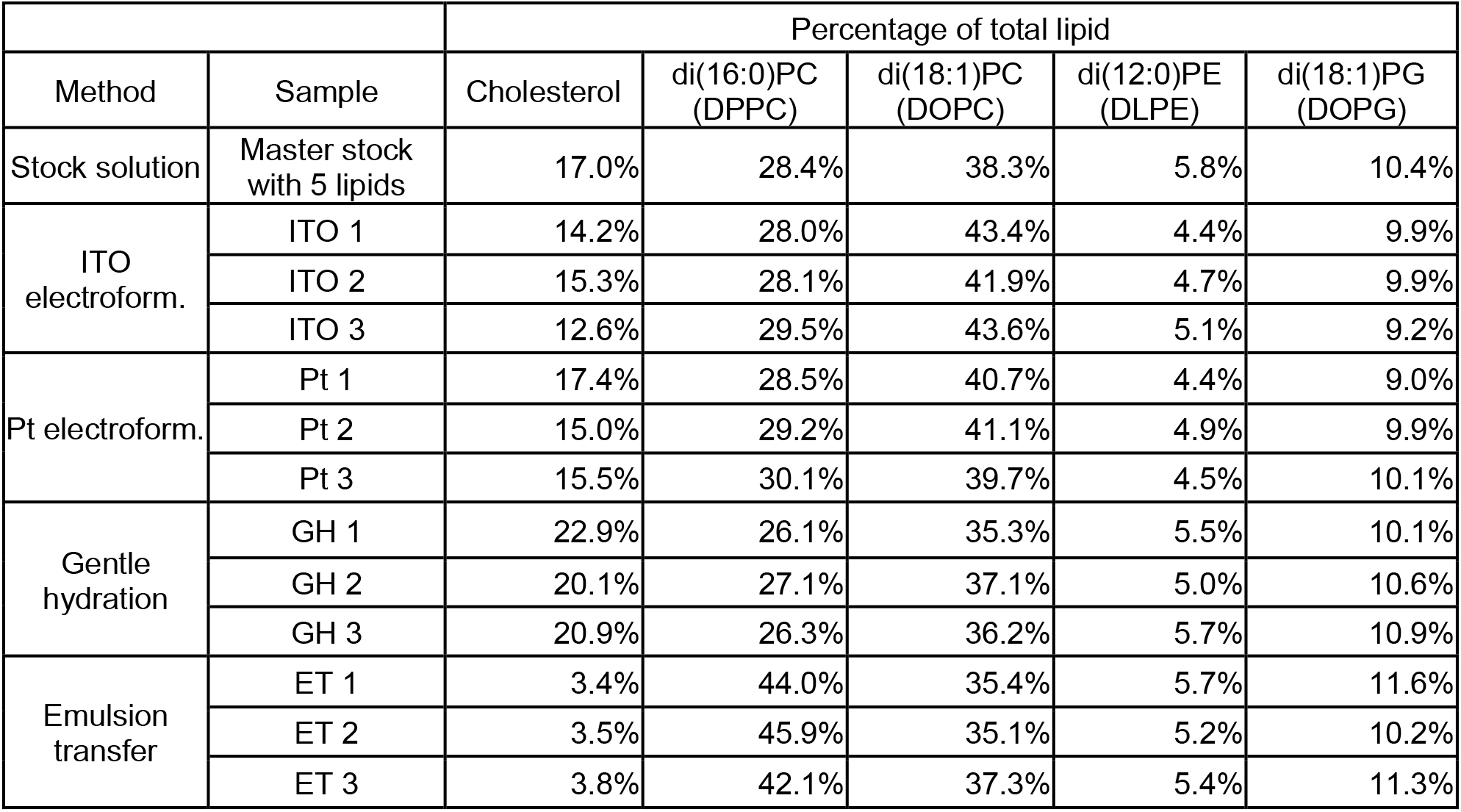
Lipid compositions of 5-component vesicles analyzed by mass spectrometry, expressed as percentages of total lipid. The abundance of each lipid is shown in Tables S11 and S12.

|  |  | Percentage of total lipid |  |  |  |  |
| --- | --- | --- | --- | --- | --- | --- |
| Method | Sample | Cholesterol | di(16:0)PC<br>(DPPC) | di(18:1)PC<br>(DOPC) | di(12:0)PE<br>(DLPE) | di(18:1)PG<br>(DOPG) |
| Stock solution | Master stock<br>with 5 lipids | 17.0% | 28.4% | 38.3% | 5.8% | 10.4% |
| ITO<br>electroform. | ITO 1 | 14.2% | 28.0% | 43.4% | 4.4% | 9.9% |
|  | ITO 2 | 15.3% | 28.1% | 41.9% | 4.7% | 9.9% |
|  | ITO 3 | 12.6% | 29.5% | 43.6% | 5.1% | 9.2% |
| Pt electroform. | Pt 1 | 17.4% | 28.5% | 40.7% | 4.4% | 9.0% |
|  | Pt 2 | 15.0% | 29.2% | 41.1% | 4.9% | 9.9% |
|  | Pt 3 | 15.5% | 30.1% | 39.7% | 4.5% | 10.1% |
| Gentle<br>hydration | GH 1 | 22.9% | 26.1% | 35.3% | 5.5% | 10.1% |
|  | GH 2 | 20.1% | 27.1% | 37.1% | 5.0% | 10.6% |
|  | GH 3 | 20.9% | 26.3% | 36.2% | 5.7% | 10.9% |
| Emulsion<br>transfer | ET 1 | 3.4% | 44.0% | 35.4% | 5.7% | 11.6% |
|  | ET 2 | 3.5% | 45.9% | 35.1% | 5.2% | 10.2% |
|  | ET 3 | 3.8% | 42.1% | 37.3% | 5.4% | 11.3% |

**Table S2.** Percentages of lipids in GUVs made by ITO electroformation, Pt wire electroformation, and emulsion transfer from a binary mixture of a saturated PC-lipid and an unsaturated PC-lipid. The abundance of each lipid is shown in Table S13. Results from an independent set of emulsion transfer experiments are shown in Tables S9 and S19.

|  |  | Percentage of total lipid |  |
| --- | --- | --- | --- |
| Method | Sample | di(16:0)PC | di(16:1)PC |
| Stock solution | Stock of saturated and unsaturated lipids in chloroform | 49.8% | 50.2% |
| ITO electroformation | ITO 1 | 46.9% | 53.1% |
|  | ITO 2 | 54.2% | 45.8% |
|  | ITO 3 | 46.6% | 53.4% |
| Pt electroformation | Pt 1 | 49.7% | 50.3% |
|  | Pt 2 | 49.0% | 51.0% |
|  | Pt 3 | 49.4% | 50.6% |
| Emulsion transfer | ET 1 | 53.1% | 46.9% |
|  | ET 2 | 60.2% | 39.8% |
|  | ET 3 | 63.2% | 36.8% |

**Table S3.**
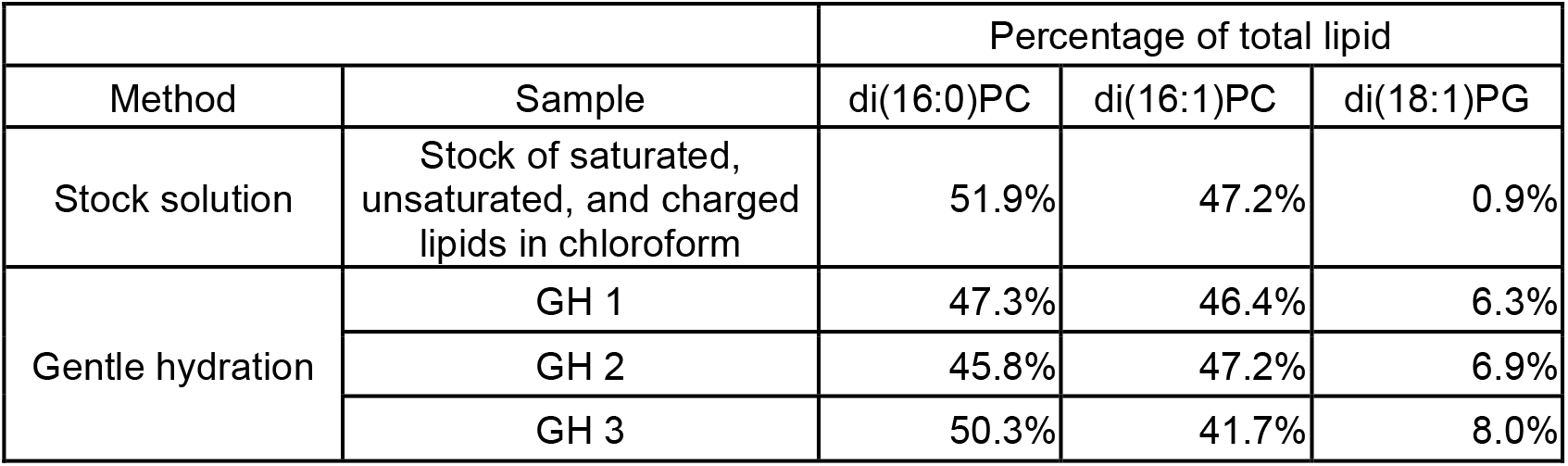
Percentages of lipids in GUVs made by gentle hydration from a mixture of a saturated and an unsaturated PC-lipid, including a charged lipid (di(18:1)PG), which is necessary for vesicle formation by this method. The abundance of each lipid is shown in Table S13.

|  |  | Percentage of total lipid |  |  |
| --- | --- | --- | --- | --- |
| Method | Sample | di(16:0)PC | di(16:1)PC | di(18:1)PG |
| Stock solution | Stock of saturated, unsaturated, and charged lipids in chloroform | 51.9% | 47.2% | 0.9% |
| Gentle hydration | GH 1 | 47.3% | 46.4% | 6.3% |
|  | GH 2 | 45.8% | 47.2% | 6.9% |
|  | GH 3 | 50.3% | 41.7% | 8.0% |

**Table S4.** Percentages of lipids in GUVs made by ITO electroformation, Pt wire electroformation, and emulsion transfer from a binary mixture of a shorter-chain PC-lipid and a longer-chain PC-lipid. The abundance of each lipid is shown in Table S14. Results from an independent set of emulsion transfer experiments are shown in Tables S10 and S20.

|  |  | Percentage of total lipid |  |
| --- | --- | --- | --- |
| Method | Sample | di(16:1)PC | di(18:1)PC |
| Stock solution | Stock of lipids with different chain lengths in chloroform | 53.4% | 46.6% |
| ITO electroformation | ITO 1 | 56.2% | 43.8% |
|  | ITO 2 | 55.9% | 44.1% |
|  | ITO 3 | 53.4% | 46.6% |
| Pt electroformation | Pt 1 | 49.7% | 50.3% |
|  | Pt 2 | 47.8% | 52.2% |
|  | Pt 3 | 46.6% | 53.4% |
| Emulsion transfer | ET 1 | 47.5% | 52.5% |
|  | ET 2 | 54.8% | 45.2% |
|  | ET 3 | 52.0% | 48.0% |

**Table S5.**
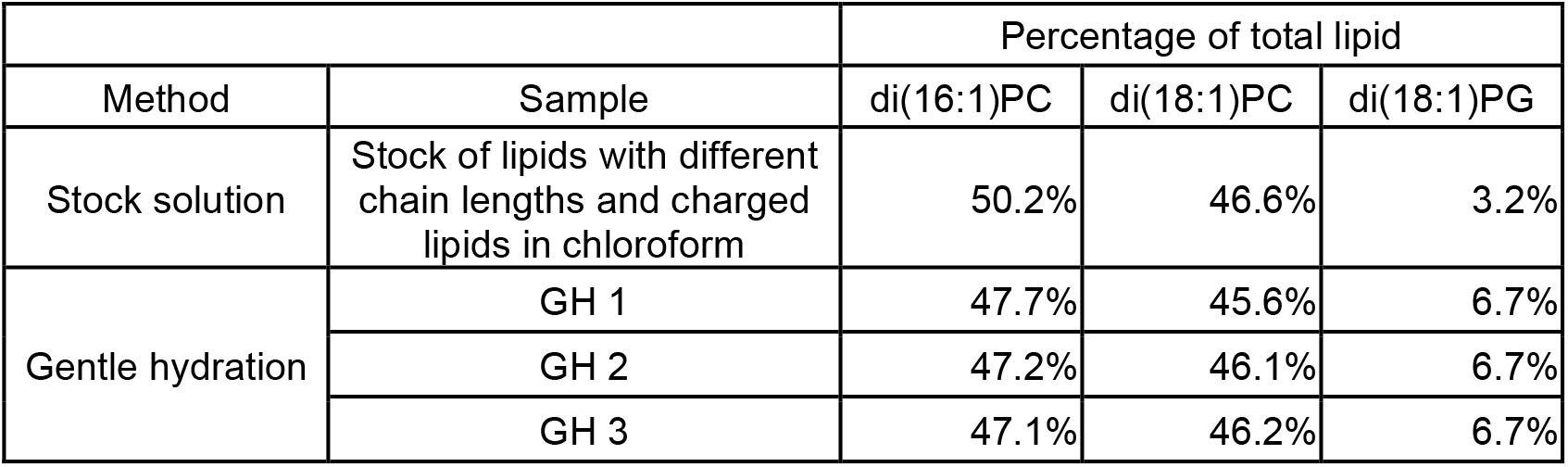
Percentages of lipids in GUVs made by gentle hydration from a mixture of a shorter-chain PC-lipid and a longer-chain PC-lipid, including a charged lipid (di(18:1)PG), which is necessary for vesicle formation by this method. The abundance of each lipid is shown in Table S14.

|  |  | Percentage of total lipid |  |  |
| --- | --- | --- | --- | --- |
| Method | Sample | di(16:1)PC | di(18:1)PC | di(18:1)PG |
| Stock solution | Stock of lipids with different chain lengths and charged lipids in chloroform | 50.2% | 46.6% | 3.2% |
| Gentle hydration | GH 1 | 47.7% | 45.6% | 6.7% |
|  | GH 2 | 47.2% | 46.1% | 6.7% |
|  | GH 3 | 47.1% | 46.2% | 6.7% |

**Table S6.** Percentages of lipids in GUVs made by ITO electroformation, Pt wire electroformation, and emulsion transfer from a binary mixture of an unsaturated PC-lipid and cholesterol. The abundance of each lipid is shown in Tables S15 and S16.

|  |  | Percentage of total lipid |  |
| --- | --- | --- | --- |
| Method | Sample | di(16:1)PC | Cholesterol |
| Stock solution | Stock of a PC-lipid and cholesterol in chloroform | 46.4% | 53.6% |
| ITO electroformation | ITO 1 | 47.1% | 52.9% |
|  | ITO 2 | 52.7% | 47.3% |
|  | ITO 3 | 47.8% | 52.2% |
| Pt electroformation | Pt 1 | 47.2% | 52.8% |
|  | Pt 2 | 47.8% | 52.2% |
|  | Pt 3 | 45.4% | 54.6% |
| Emulsion transfer | ET 1 | 80.3% | 19.7% |
|  | ET 2 | 92.9% | 7.1% |
|  | ET 3 | 87.8% | 12.2% |

**Table S7.** Percentages of lipids in GUVs made by gentle hydration from a mixture of an unsaturated PC-lipid and cholesterol, including a charged lipid (di(18:1)PG), which is necessary for vesicle formation by this method. The abundance of each lipid is shown in Tables S15 and S16.

|  |  | Percentage of total lipid |  |  |
| --- | --- | --- | --- | --- |
| Method | Sample | di(16:1)PC | Cholesterol | di(18:1)PG |
| Stock solution | Stock of a PC-lipid, cholesterol, and a charged lipid in chloroform | 41.6% | 56.8% | 1.6% |
| Gentle hydration | GH 1 | 29.8% | 66.8% | 3.4% |
|  | GH 2 | 36.5% | 58.4% | 5.2% |
|  | GH 3 | 33.0% | 62.7% | 4.3% |

**Table S8.** Lipid compositions analyzed by mass spectrometry and expressed as percentages of total lipid for vesicles produced by gentle hydration (“before extrusion”) and then extruded (“after extrusion”). For example, “Experiment 1 – After" was made from an aliquot of the solution from “Experiment 1 – Before". The abundance of each lipid is shown in Tables S17 and S18.

| Method | Sample | Percentage of total lipid |  |  |  |  |
| --- | --- | --- | --- | --- | --- | --- |
|  |  | Cholesterol | di(16:0)PC (DPPC) | di(18:1)PC (DOPC) | di(12:0)PE (DLPE) | di(18:1)PG (DOPG) |
| Before extrusion | Experiment 1 - Before | 23.6% | 25.2% | 29.2% | 6.9% | 15.0% |
|  | Experiment 2 - Before | 26.3% | 26.5% | 25.6% | 6.5% | 15.1% |
|  | Experiment 3 - Before | 27.7% | 25.8% | 25.2% | 6.3% | 14.9% |
| After extrusion | Experiment 1 - After | 18.0% | 27.9% | 32.1% | 5.2% | 16.9% |
|  | Experiment 2 - After | 22.4% | 27.3% | 27.1% | 6.7% | 16.6% |
|  | Experiment 3 - After | 22.3% | 27.0% | 28.4% | 6.7% | 15.6% |

**Table S9.** Percentages of lipids in GUVs made by emulsion transfer from an additional binary mixture of a saturated PC-lipid and an unsaturated PC-lipid. The abundance of each lipid is shown in Table S19. Data from an independent set of emulsion transfer experiments are shown in Tables S2 and S13.

| Method | Sample | Percentage of total lipid |  |
| --- | --- | --- | --- |
|  |  | di(16:0)PC | di(16:1)PC |
| Stock solution for lipid saturation experiments | Stock of saturated and unsaturated lipids in chloroform | 49.1% | 50.9% |
| Emulsion transfer (ET) | ET saturation 1 | 58.2% | 41.8% |
|  | ET saturation 2 | 58.0% | 42.0% |
|  | ET saturation 3 | 57.3% | 42.7% |

**Table S10.** Percentages of lipids in GUVs made by emulsion transfer from an additional binary mixture of a shorter-chain PC-lipid and a longer-chain PC-lipid. The abundance of each lipid is shown in Table S20. Data from an independent set of emulsion transfer experiments are shown in Tables S4 and S14.

| Method | Sample | Percentage of total lipid |  |
| --- | --- | --- | --- |
|  |  | di(18:1)PC | di(16:1)PC |
| Stock solution for lipid length experiments | Stock of lipids with different chain lengths in chloroform | 49.1% | 50.9% |
| Emulsion transfer (ET) | ET length 1 | 58.8% | 41.2% |
|  | ET length 2 | 58.7% | 41.3% |
|  | ET length 3 | 53.1% | 46.9% |

**Table S11.** Phospholipid abundances of 5-component vesicles analyzed by mass spectrometry. Corresponding lipid percentages are in Table S1.

| Lipid | di(15:0)PC<br>*Internal*<br>*standard* | di(16:0)PC | di(18:1)PC | di(15:0)PE<br>*Internal*<br>*standard* | di(12:0)PE | di(15:0)PG<br>*Internal*<br>*standard* | di(18:1)PG |
| --- | --- | --- | --- | --- | --- | --- | --- |
| m/z | 706.5 | 734.6 | 786.6 | 686.5 | 602.4 | 717.5 | 797.5 |
| Retention time<br>(minutes) | 3.467 | 3.467 | 3.414 | 2.725 | 2.847 | 0.708 | 0.621 |
| Calibrated retention<br>time (minutes) | 6.760 | 6.760 | 6.672 | 5.542 | 5.742 | 2.233 | 2.038 |
| Collisional cross<br>section (Å<sup>2</sup>) | 274.4 | 279.8 | 285.1 | 268.9 | 252.5 | 268.6 | 279.4 |
| 5-component stock<br>solution | 5617.5 | 17245.3 | 23270.3 | 4528.2 | 2853.3 | 4441.4 | 5016.5 |
| ITO electroform. 1<br>abundance | 6741.8 | 17442.6 | 27025.4 | 4434.9 | 1810.2 | 5385.0 | 4914.3 |
| ITO electroform. 2<br>abundance | 6857.4 | 24043.3 | 35779.5 | 4280.0 | 2523.2 | 5170.9 | 6393.5 |
| ITO electroform. 3<br>abundance | 7240.7 | 15652.1 | 23118.4 | 4920.6 | 1843.7 | 6871.4 | 4625.1 |
| Pt electroform. 1<br>abundance | 6881.0 | 22694.3 | 32436.7 | 4665.2 | 2379.2 | 5772.2 | 6026.1 |
| Pt electroform. 2<br>abundance | 6679.0 | 16801.5 | 23631.0 | 5039.4 | 2124.3 | 6748.9 | 5740.9 |
| Pt electroform. 3<br>abundance | 5342.4 | 14477.7 | 19085.5 | 4548.3 | 1847.1 | 5020.5 | 4557.4 |
| Gentle hydration 1<br>abundance | 7353.0 | 30264.4 | 40917.1 | 5487.3 | 4792.2 | 5907.4 | 9413.3 |
| Gentle hydration 2<br>abundance | 7682.1 | 23456.5 | 32112.9 | 6630.8 | 3748.8 | 5247.5 | 6246.9 |
| Gentle hydration 3<br>abundance | 7625.5 | 22749.8 | 31393.1 | 5379.8 | 3504.5 | 5017.3 | 6195.7 |
| Emulsion transfer 1<br>abundance | 8370.8 | 9579.3 | 7712.5 | 5534.3 | 816.3 | 6091.2 | 1843.4 |
| Emulsion transfer 2<br>abundance | 8068.4 | 11843.5 | 9051.1 | 5553.9 | 931.3 | 5927.0 | 1931.0 |
| Emulsion transfer 3<br>abundance | 7626.0 | 9787.1 | 8667.5 | 5246.7 | 867.2 | 5531.6 | 1910.8 |

**Table S12.** Cholesterol abundances of 5-component vesicles analyzed by mass spectrometry. Corresponding phospholipid abundances are in Table S11. Corresponding lipid percentages are in Table S1. “CholesterolStandardMix” samples contained 1 µg/mL of d7-cholesterol and 0.2 µg/mL of unlabeled cholesterol and were used to calculate an RRF value (right column). For example, the RRF for CholesterolStandardMix1 is calculated: (0.2 µg/mL / 3,700,000) / (1 µg/mL / 19,200,000) = 1.038. The average RRF value 1.063 was used to calculate the cholesterol composition in all 5-component experimental samples.

|  | d7-cholesterol<br>*Internal standard* | Cholesterol | Relative Response<br>Factor (RRF) |
| --- | --- | --- | --- |
| 5-component stock solution | 321. x 10<sup>5</sup> | 85.9 x 10<sup>5</sup> | 1.063 |
| ITO electroformation 1 | 268. x 10<sup>5</sup> | 51.3 x 10<sup>5</sup> |  |
| ITO electroformation 2 | 276. x 10<sup>5</sup> | 76.7 x 10<sup>5</sup> |  |
| ITO electroformation 3 | 253. x 10<sup>5</sup> | 33.9 x 10<sup>5</sup> |  |
| Pt electroformation 1 | 284. x 10<sup>5</sup> | 83.3 x 10<sup>5</sup> |  |
| Pt electroformation 2 | 261. x 10<sup>5</sup> | 49.1 x 10<sup>5</sup> |  |
| Pt electroformation 3 | 262. x 10<sup>5</sup> | 53.3 x 10<sup>5</sup> |  |
| Gentle hydration 1 | 299. x 10<sup>5</sup> | 157. x 10<sup>5</sup> |  |
| Gentle hydration 2 | 271. x 10<sup>5</sup> | 89.3 x 10<sup>5</sup> |  |
| Gentle hydration 3 | 233. x 10<sup>5</sup> | 80.5 x 10<sup>5</sup> |  |
| Emulsion transfer 1 | 223. x 10<sup>5</sup> | 2.85 x 10<sup>5</sup> |  |
| Emulsion transfer 2 | 235. x 10<sup>5</sup> | 3.88 x 10<sup>5</sup> |  |
| Emulsion transfer 3 | 367. x 10<sup>5</sup> | 6.13 x 10<sup>5</sup> |  |
| CholesterolStandardMix 1 | 192. x 10<sup>5</sup> | 37.0 x 10<sup>5</sup> | 1.038 |
| CholesterolStandardMix 2 | 195. x 10<sup>5</sup> | 37.5 x 10<sup>5</sup> | 1.040 |
| CholesterolStandardMix 3 | 192. x 10<sup>5</sup> | 36.4 x 10<sup>5</sup> | 1.055 |
| CholesterolStandardMix 4 | 174. x 10<sup>5</sup> | 33.1 x 10<sup>5</sup> | 1.051 |
| CholesterolStandardMix 5 | 184. x 10<sup>5</sup> | 32.6 x 10<sup>5</sup> | 1.129 |

**Table S13.** Phospholipid abundances in a stock solution and in giant unilamellar vesicles made by ITO electroformation, Pt wire electroformation, emulsion transfer and gentle hydration from a binary mixture of a saturated PC-lipid and an unsaturated PC-lipid. Samples for gentle hydration included a charged lipid (di(18:1)PG), which is necessary for vesicle formation by this method. Corresponding lipid percentages are in Tables S2 and S3.

| Lipid | di(15:0)PC<br>*Internal*<br>*standard* | di(16:0)PC | (16:0/18:1)PC<br>*Internal*<br>*standard* | di(16:1)PC | di(15:0)PG<br>*Internal*<br>*standard* | di(18:1)PG |
| --- | --- | --- | --- | --- | --- | --- |
| m/z | 706.5 | 734.6 | 760.6 | 730.5 | 717.5 | 797.5 |
| Retention time<br>(minutes) | 3.541 | 3.523 | 3.478 | 3.495 | 0.953 | 0.838 |
| Calibrated<br>retention time<br>(minutes) | 6.789 | 6.761 | 6.685 | 6.713 | 2.233 | 2.038 |
| Collisional cross<br>section ($\text{\AA}^2$) | 240.8 | 249.6 | 248.3 | 241.8 | 238.5 | 247.0 |
| Stock for lipid<br>saturation tests | 30860.0 | 43216.7 | 31604.8 | 44611.7 |  |  |
| ITO electroform. 1<br>abundance | 24366.7 | 14971.9 | 24647.1 | 17159.7 |  |  |
| ITO electroform. 2<br>abundance | 35109.7 | 48160.5 | 39166.9 | 45440.7 |  |  |
| ITO electroform. 3<br>abundance | 33117.1 | 42203.3 | 29583.7 | 43222.0 |  |  |
| Pt. electroform. 1<br>abundance | 33264.2 | 10641.9 | 35170.3 | 11384.6 |  |  |
| Pt. electroform. 2<br>abundance | 33512.3 | 8938.4 | 36105.9 | 10028.2 |  |  |
| Pt. electroform. 3<br>abundance | 27387.5 | 13675.6 | 28251.8 | 14440.8 |  |  |
| Emulsion transfer 1<br>abundance | 20453.6 | 24519.6 | 21912.5 | 23201.6 |  |  |
| Emulsion transfer 2<br>abundance | 28376.8 | 29801.8 | 35758.6 | 24827.5 |  |  |
| Emulsion transfer 3<br>abundance | 28682.9 | 10546.0 | 29830.2 | 6389.1 |  |  |
| Gentle hydration<br>stock solution for<br>lipid saturation | 24928.8 | 211952.6 | 28549.2 | 221074.0 | 50120.7 | 7402.7 |
| Gentle hydration 1<br>abundance | 22683.3 | 271061.2 | 24664.9 | 289244.2 | 37829.3 | 60299.3 |
| Gentle hydration 2<br>abundance | 19336.3 | 193665.6 | 22045.5 | 227491.3 | 26953.7 | 40829.7 |
| Gentle hydration 3<br>abundance | 24807.4 | 214717.1 | 31620.1 | 227134.8 | 35032.4 | 48250.1 |

**Table S14.**
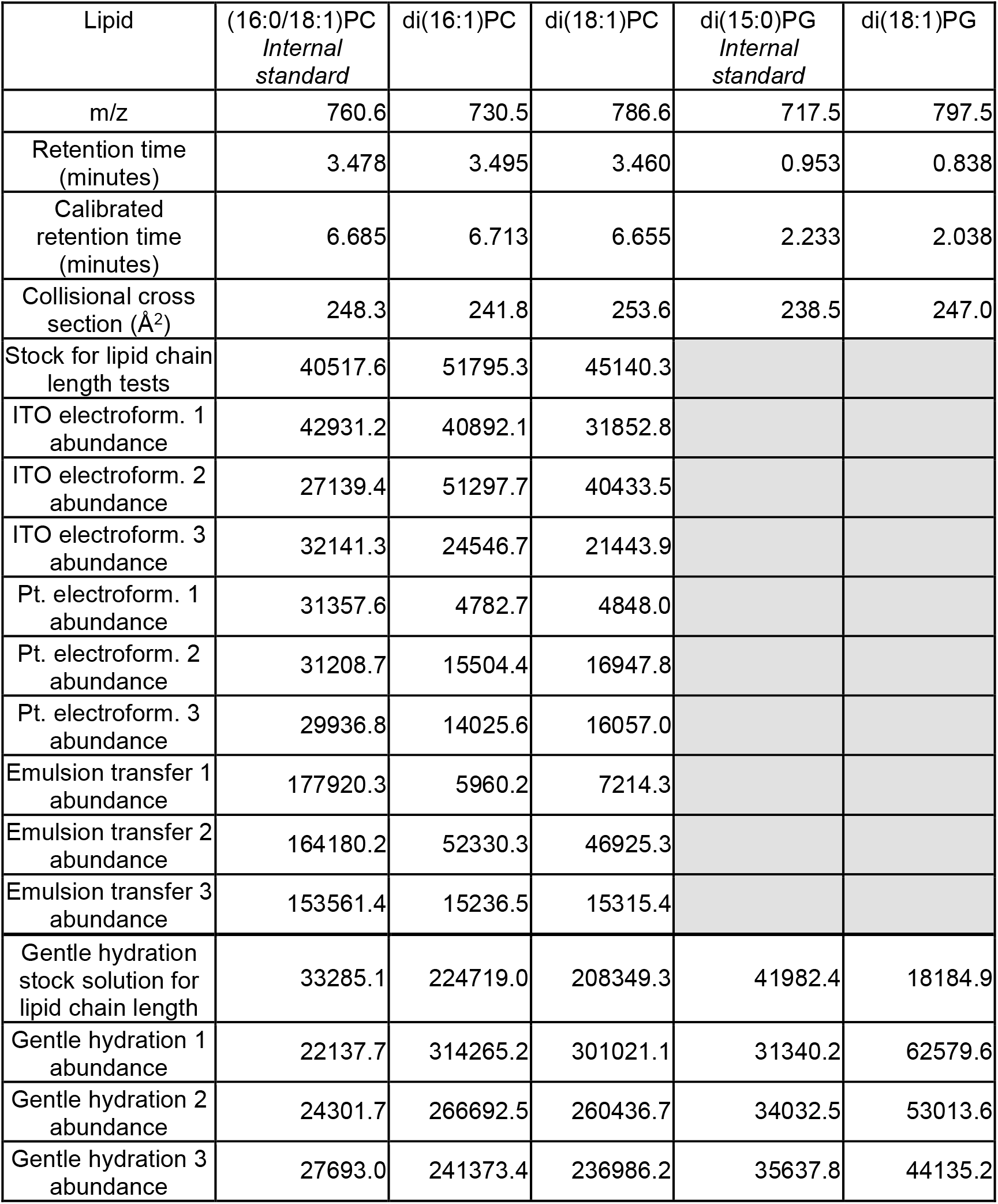
Phospholipid abundances in stock solutions and in giant unilamellar vesicles made by ITO electroformation, Pt wire electroformation, emulsion transfer, and gentle hydration from a mixture of a lipids with different chain lengths. Samples for gentle hydration included a charged lipid (di(18:1)PG), which is necessary for vesicle formation by this method. Corresponding lipid percentages are in Tables S4 and S5.

**Table S15.** Phospholipid abundances in stock solutions and in giant unilamellar vesicles made by ITO electroformation, Pt wire electroformation, emulsion transfer, and gentle hydration from a mixture of a PC-lipid and cholesterol. Samples for gentle hydration included a charged lipid (di(18:1)PG), which is necessary for vesicle formation by this method. Corresponding cholesterol abundances are in Table S16. Corresponding lipid percentages are in Tables S6 and S7.

| Lipid | (16:0)(18:1)PC<br>*Internal*<br>*standard* | di(16:1)PC | di(15:0)PG<br>*Internal*<br>*standard* | di(18:1)PG |
| --- | --- | --- | --- | --- |
| m/z | 760.6 | 730.5 | 717.5 | 797.5 |
| Retention time<br>(minutes) | 3.478 | 3.495 | 0.953 | 0.838 |
| Calibrated<br>retention time<br>(minutes) | 6.685 | 6.713 | 2.233 | 2.038 |
| Collisional cross<br>section ($\text{\AA}^2$) | 248.3 | 241.8 | 238.5 | 247.0 |
| Stock for PC-lipid<br>vs. cholesterol | 31657.8 | 46654.6 |  |  |
| ITO electroform. 1<br>abundance | 30214.6 | 35616.8 |  |  |
| ITO electroform. 2<br>abundance | 26795.1 | 54201.0 |  |  |
| ITO electroform. 3<br>abundance | 36225.0 | 59824.3 |  |  |
| Pt. electroform. 1<br>abundance | 32660.1 | 20906.6 |  |  |
| Pt. electroform. 2<br>abundance | 34421.4 | 40375.1 |  |  |
| Pt. electroform. 3<br>abundance | 33803.4 | 31500.6 |  |  |
| Emulsion transfer 1<br>abundance | 162535.6 | 41325.6 |  |  |
| Emulsion transfer 2<br>abundance | 152545.4 | 195281.5 |  |  |
| Emulsion transfer 3<br>abundance | 159797.7 | 90395.7 |  |  |
| Gentle hydration<br>stock for PC-lipid<br>vs. cholesterol | 33090.1 | 272405.7 | 47084.9 | 14735.5 |
| Gentle hydration 1<br>abundance | 23214.1 | 341451.9 | 32808.2 | 54979.6 |
| Gentle hydration 2<br>abundance | 24085.7 | 296331.5 | 30982.6 | 53893.9 |
| Gentle hydration 3<br>abundance | 26059.0 | 306777.7 | 30714.2 | 47246.0 |

**Table S16.** Cholesterol abundances in stock solutions and in giant unilamellar vesicles made by ITO electroformation, Pt wire electroformation, emulsion transfer, and gentle hydration from a mixture of a PC-lipid and cholesterol. Corresponding phospholipid abundances are in Table S15. Corresponding lipid percentages are in Tables S6 and S7. “Cholesterol Standard Mix” samples contained 1 µg/mL of d7-cholesterol and 0.2 µg/mL of unlabeled cholesterol and were used to calculate an RRF value (right column). For example, the RRF for Cholesterol Standard Mix 1 is calculated: (0.2 µg/mL / 3297000) / (1 µg/mL / 22310000) = 0.7389. Cholesterol Standard Mix 1-9 were used to calculate an average RRF value 0.7511 which was used to calculate the cholesterol compositions for all binary PC-lipid vs. cholesterol experimental samples, excluding emulsion transfer. For emulsion transfer, Cholesterol Standard Mix 10-13 were used to calculate an average RRF value 0.8300 which was used to calculate cholesterol compositions after the samples were concentrated by a factor of 5.

|  | d7-cholesterol<br>*Internal standard* | Cholesterol | Relative Response<br>Factor (RRF) |
| --- | --- | --- | --- |
| Stock for PC-lipid vs.<br>cholesterol | 71.91 x 10<sup>5</sup> | 13.64 x 10<sup>5</sup> | 0.7511 |
| ITO electroformation 1 | 127.5 x 10<sup>5</sup> | 18.31 x 10<sup>5</sup> |  |
| ITO electroformation 2 | 116.1 x 10<sup>5</sup> | 23.90 x 10<sup>5</sup> |  |
| ITO electroformation 3 | 154.8 x 10<sup>5</sup> | 30.65 x 10<sup>5</sup> |  |
| Pt electroform 1 | 154.4 x 10<sup>5</sup> | 9.520 x 10<sup>5</sup> |  |
| Pt electroform 2 | 158.9 x 10<sup>5</sup> | 20.60 x 10<sup>5</sup> |  |
| Pt electroform 3 | 160.3 x 10<sup>5</sup> | 17.96 x 10<sup>5</sup> |  |
| Emulsion transfer 1 | 1019. x 10<sup>5</sup> | 9.249 x 10<sup>5</sup> | 0.8300 |
| Emulsion transfer 2 | 308.5 x 10<sup>5</sup> | 4.048 x 10<sup>5</sup> |  |
| Emulsion transfer 3 | 18.63 x 10<sup>5</sup> | 0.2113 x 10<sup>5</sup> |  |
| Gentle hydration stock for PC-lipid vs. cholesterol | 148.7 x 10<sup>5</sup> | 211.9 x 10<sup>5</sup> | 0.7511 |
| Gentle hydration 1 | 169.4 x 10<sup>5</sup> | 701.6 x 10<sup>5</sup> |  |
| Gentle hydration 2 | 163.2 x 10<sup>5</sup> | 406.1 x 10<sup>5</sup> |  |
| Gentle hydration 3 | 160.1 x 10<sup>5</sup> | 454.3 x 10<sup>5</sup> |  |
| Cholesterol Standard Mix 1 | 223.1 x 10<sup>5</sup> | 32.97 x 10<sup>5</sup> | 0.7389 |
| Cholesterol Standard Mix 2 | 224.1 x 10<sup>5</sup> | 32.89 x 10<sup>5</sup> | 0.7339 |
| Cholesterol Standard Mix 3 | 221.1 x 10<sup>5</sup> | 32.76 x 10<sup>5</sup> | 0.7411 |
| Cholesterol Standard Mix 4 | 230.0 x 10<sup>5</sup> | 34.46 x 10<sup>5</sup> | 0.7492 |
| Cholesterol Standard Mix 5 | 225.5 x 10<sup>5</sup> | 34.06 x 10<sup>5</sup> | 0.7553 |
| Cholesterol Standard Mix 6 | 224.7 x 10<sup>5</sup> | 34.70 x 10<sup>5</sup> | 0.7721 |
| Cholesterol Standard Mix 7 | 231.6 x 10<sup>5</sup> | 35.56 x 10<sup>5</sup> | 0.7677 |
| Cholesterol Standard Mix 8 | 237.2 x 10<sup>5</sup> | 35.49 x 10<sup>5</sup> | 0.7480 |
| Cholesterol Standard Mix 9 | 240.5 x 10<sup>5</sup> | 36.25 x 10<sup>5</sup> | 0.7536 |
| Cholesterol Standard Mix 10 | 347.6 x 10<sup>5</sup> | 57.44 x 10<sup>5</sup> | 0.8263 |
| Cholesterol Standard Mix 11 | 342.1 x 10<sup>5</sup> | 57.29 x 10<sup>5</sup> | 0.8373 |
| Cholesterol Standard Mix 12 | 335.9 x 10<sup>5</sup> | 55.55 x 10<sup>5</sup> | 0.8268 |
| Cholesterol Standard Mix 13 | 331.1 x 10<sup>5</sup> | 54.95 x 10<sup>5</sup> | 0.8297 |

**Table S17.** Phospholipid abundances in giant unilamellar vesicles produced by gentle hydration (“before extrusion”) and then extruded (“after extrusion”). For example, “After extrusion 1” was made from an aliquot of the solution from “Before extrusion 1”. Corresponding lipid percentages are in Table S8.

| Lipid | di(15:0)PC<br>*Internal*<br>*standard* | di(16:0)PC | di(18:1)PC | di(15:0)PE<br>*Internal*<br>*standard* | di(12:0)PE | di(15:0)PG<br>*Internal*<br>*standard* | di(18:1)PG |
| --- | --- | --- | --- | --- | --- | --- | --- |
| m/z | 706.5 | 734.6 | 786.6 | 664.5 | 580.4 | 717.5 | 797.5 |
| Retention time<br>(minutes) | 3.312 | 3.312 | 3.225 | 2.570 | 2.692 | 0.674 | 0.553 |
| Calibrated retention<br>time (minutes) | 5.445 | 5.445 | 5.459 | 5.560 | 5.541 | 2.233 | 2.020 |
| Collisional cross<br>section (Å<sup>2</sup>) | 239.9 | 249.9 | 254.0 | 232.0 | 216.8 | 239.8 | 248.4 |
| Before extrusion 1<br>abundance | 56822.3 | 84583.4 | 97994.1 | 85471.0 | 34719.6 | 36543.5 | 32416.2 |
| Before extrusion 2<br>abundance | 63191.0 | 113815.1 | 109639.1 | 101636.3 | 44984.7 | 39897.5 | 40816.1 |
| Before extrusion 3<br>abundance | 63430.2 | 159911.8 | 156115.9 | 95764.3 | 59014.5 | 30611.1 | 44526.0 |
| After extrusion 1<br>abundance | 54683.8 | 32544.4 | 37473.0 | 89296.3 | 9911.4 | 43706.7 | 15788.9 |
| After extrusion 2<br>abundance | 65041.1 | 65510.5 | 65082.1 | 96925.9 | 23877.9 | 37763.1 | 23185.0 |
| After extrusion 3<br>abundance | 50216.3 | 63415.0 | 66752.6 | 85189.5 | 26663.7 | 36801.5 | 26753.6 |

**Table S18.** Cholesterol abundances in giant unilamellar vesicles produced by gentle hydration (“before extrusion”) and then extruded (“after extrusion”). For example, “After extrusion 1” was made from an aliquot of the solution from “Before extrusion 1”. Corresponding phospholipid abundances are in Table S17. Corresponding lipid percentages are in Table S8. “CholesterolStandardMix” samples contained 1 µg/mL of d7-cholesterol and 0.2 µg/mL of unlabeled cholesterol and were used to calculate an RRF value (right column). For example, the RRF for CholesterolStandardMix1 is calculated: (0.2 µg/mL / 4,244,000) / (1 µg/mL / 18,350,000) = 0.8650. The average RRF value 0.8608 was used to calculate the cholesterol composition in all 5-component experimental samples.

| Sample | d7-cholesterol/*Internal standard* | Cholesterol | Relative Response Factor (RRF) |
| --- | --- | --- | --- |
| 5-component stock solution | 161.0 x 10<sup>5</sup> | 3879. x 10<sup>5</sup> | 0.8608 |
| Before extrusion 1 | 170.1 x 10<sup>5</sup> | 42.56 x 10<sup>5</sup> |  |
| Before extrusion 2 | 169.5 x 10<sup>5</sup> | 54.41 x 10<sup>5</sup> |  |
| Before extrusion 3 | 165.1 x 10<sup>5</sup> | 80.23 x 10<sup>5</sup> |  |
| After extrusion 1 | 158.4 x 10<sup>5</sup> | 10.93 x 10<sup>5</sup> |  |
| After extrusion 2 | 166.2 x 10<sup>5</sup> | 24.68 x 10<sup>5</sup> |  |
| After extrusion 3 | 154.1 x 10<sup>5</sup> | 28.86 x 10<sup>5</sup> |  |
| CholesterolStandardMix 1 | 183.5 x 10<sup>5</sup> | 42.44 x 10<sup>5</sup> | 0.8650 |
| CholesterolStandardMix 2 | 185.1 x 10<sup>5</sup> | 43.33 x 10<sup>5</sup> | 0.8545 |
| CholesterolStandardMix 3 | 182.6 x 10<sup>5</sup> | 42.33 x 10<sup>5</sup> | 0.8628 |

**Table S19.** Phospholipid abundances in giant unilamellar vesicles made for an additional emulsion transfer triplicate from a binary mixture of an unsaturated lipid and a saturated lipid. Corresponding lipid percentages are in Table S9. Data from an independent set of emulsion transfer experiments are shown in Tables S2 and S13.

| Lipid | di(15:0)PC<br>*Internal standard* | di(16:0)PC | (16:0/18:1)PC<br>*Internal standard* | di(16:1)PC |
| --- | --- | --- | --- | --- |
| m/z | 706.5 | 734.6 | 760.6 | 730.5 |
| Retention time (minutes) | 3.312 | 3.312 | 3.259 | 3.414 |
| Calibrated retention time (minutes) | 5.445 | 5.445 | 5.453 | 5.429 |
| Collisional cross section (Å<sup>2</sup>) | 239.9 | 249.9 | 248.7 | 240.9 |
| Emulsion transfer stock for lipid saturation tests | 14259.5 | 490722.3 | 12208.1 | 434934.4 |
| ET saturation 1 abundance | 54941.2 | 45862.2 | 52621.1 | 31490.8 |
| ET saturation 2 abundance | 58862.0 | 21123.8 | 55943.2 | 14554.5 |
| ET saturation 3 abundance | 57614.3 | 39973.9 | 54335.7 | 28099.2 |

**Table S20.** Phospholipid abundances in GUVs made for an additional emulsion transfer triplicate from a binary mixture of lipids with different chain lengths. Corresponding lipid percentages are in Table S10. Data from an independent set of emulsion transfer experiments are shown in Tables S4 and S14.

| Lipid | (16:0/18:1)PC<br>*Internal standard* | di(16:1)PC | di(18:1)PC |
| --- | --- | --- | --- |
| m/z | 760.6 | 730.5 | 786.6 |
| Retention time (minutes) | 3.259 | 3.414 | 3.225 |
| Calibrated retention time (minutes) | 5.453 | 5.429 | 5.459 |
| Collisional cross section (Å<sup>2</sup>) | 248.7 | 240.9 | 254.0 |
| Emulsion transfer stock for lipid chain length tests | 8362.4 | 428416.6 | 412967.2 |
| ET length 1 abundance | 51197.1 | 13380.6 | 19090.4 |
| ET length 2 abundance | 53042.6 | 5718.8 | 8128.1 |
| ET length 3 abundance | 54721.9 | 10486.8 | 11892.0 |

## REFERENCES

1. Olson, F., C.A. Hunt, F.C. Szoka, W.J. Vail, and D. Papahadjopoulos. 1979. Preparation of liposomes of defined size distribution by extrusion through polycarbonate membranes. BBA - Biomembr. 557:9–23.

2. Sugiura, S., T. Kuroiwa, T. Kagota, M. Nakajima, S. Sato, S. Mukataka, P. Walde, and S. Ichikawa. 2008. Novel Method for Obtaining Homogeneous Giant Vesicles from a Monodisperse Water-in-Oil Emulsion Prepared with a Microfluidic Device. Langmuir. 24:4581–4588.

3. Zhu, T.F., and J.W. Szostak. 2009. Preparation of Large Monodisperse Vesicles. PLoS ONE. 4:e5009.

4. Angelova, M.I. 1988. Lipid Swelling and Liposome Formation in Electric Fields. PhD thesis. Bulgarian Academy of Sciences.

5. Ghellab, S.E., Q. Li, T. Fuhs, H. Bi, and X. Han. 2017. Electroformation of double vesicles using an amplitude modulated electric field. Colloids Surf. B Biointerfaces. 160:697–703.

6. Ip, T., Q. Li, N. Brooks, and Y. Elani. 2021. Manufacture of Multilayered Artificial Cell Membranes through Sequential Bilayer Deposition on Emulsion Templates. ChemBioChem. 22:2275–2281.

7. Pautot, S., B.J. Frisken, and D.A. Weitz. 2003. Engineering asymmetric vesicles. Proc. Natl. Acad. Sci. 100:10718–10721.

8. Elani, Y., S. Purushothaman, P.J. Booth, J.M. Seddon, N.J. Brooks, R.V. Law, and O. Ces. 2015. Measurements of the effect of membrane asymmetry on the mechanical properties of lipid bilayers. Chem. Commun. 51:6976–6979.

9. Doktorova, M., F.A. Heberle, B. Eicher, R.F. Standaert, J. Katsaras, E. London, G. Pabst, and D. Marquardt. 2018. Preparation of asymmetric phospholipid vesicles: The next generation of cell membrane models. Nat. Protoc. 13:2086–2101.

10. Akashi, K., H. Miyata, H. Itoh, and K. Kinosita. 1996. Preparation of giant liposomes in physiological conditions and their characterization under an optical microscope. Biophys. J. 71:3242–3250.

11. Blosser, M.C., J.B. Starr, C.W. Turtle, J. Ashcraft, and S.L. Keller. 2013. Minimal Effect of Lipid Charge on Membrane Miscibility Phase Behavior in Three Ternary Systems. Biophys. J. 104:2629–2638.

12. Dominak, L.M., and C.D. Keating. 2007. Polymer Encapsulation within Giant Lipid Vesicles. Langmuir. 23:7148–7154.

13. Abkarian, M., E. Loiseau, and G. Massiera. 2011. Continuous droplet interface crossing encapsulation (cDICE) for high throughput monodisperse vesicle design. Soft Matter. 7:4610–4614.

14. Dimova, R., P. Stano, C. M. Marques, P. Walde, 2019. The Giant Vesicle Book, Chapter 1. Preparation methods for giant unilamellar vesicles. R. Dimova and C. M. Marques, editors. CRC Press, Boca Raton, pp. 1–16.

15. Huang, J., J.T. Buboltz, and G.W. Feigenson. 1999. Maximum solubility of cholesterol in phosphatidylcholine and phosphatidylethanolamine bilayers. BBA - Biomembr. 1417:89– 100.

16. Epand, R.M., D.W. Hughes, B.G. Sayer, N. Borochov, D. Bach, and E. Wachtel. 2003. Novel properties of cholesterol-dioleoylphosphatidylcholine mixtures. Biochim. Biophys. Acta. 1616:196–208.

17. Shaikh, S.R., V. Cherezov, M. Caffrey, S.P. Soni, D. LoCascio, W. Stillwell, and S.R. Wassall. 2006. Molecular Organization of Cholesterol in Unsaturated Phosphatidylethanolamines: X-ray Diffraction and Solid State 2H NMR Reveal Differences with Phosphatidylcholines. J. Am. Chem. Soc. 128:5375–5383.

18. Stevens, M.M., A.R. Honerkamp-Smith, and S.L. Keller. 2010. Solubility Limits of Cholesterol, Lanosterol, Ergosterol, Stigmasterol, and β-Sitosterol in Electroformed Lipid Vesicles. Soft Matter. 6:5882–5890.

19. Ibarguren, M., A. Alonso, B.G. Tenchov, and F.M. Goñi. 2010. Quantitation of cholesterol incorporation into extruded lipid bilayers. BBA - Biomembr. 1798:1735–1738.

20. Blosser, M.C., B.G. Horst, and S.L. Keller. 2016. cDICE method produces giant lipid vesicles under physiological conditions of charged lipids and ionic solutions. Soft Matter. 12:7364–7371.

21. Karamdad, K., J. W. Hindley, G. Bolognesi, M. S. Friddin, R. V. Law, N. J. Brooks, O. Ces, and Y. Elani. 2018. Engineering thermoresponsive phase separated vesicles formed via emulsion phase transfer as a content-release platform. Chem. Sci. 9:4851–4858.

22. Dürre, K., and A. R. Bausch. 2019. Formation of phase separated vesicles by double layer cDICE. Soft Matter. 15:9676–9681.

23. Veatch, S.L., and S.L. Keller. 2005. Seeing spots: Complex phase behavior in simple membranes. BBA - Mol. Cell Res. 1746:172–185.

24. Gambhir, A., G. Hangyás-Mihályné, I. Zaitseva, D.S. Cafiso, J. Wang, D. Murray, S.N. Pentyala, S.O. Smith, and S. McLaughlin. 2004. Electrostatic Sequestration of PIP2 on Phospholipid Membranes by Basic/Aromatic Regions of Proteins. Biophys. J. 86:2188– 2207.

25. Cullis, P.R., M.J. Hope, and C.P.S. Tilcock. 1986. Lipid polymorphism and the roles of lipids in membranes. Chem. Phys. Lipids. 40:127–144.

26. van der Veen, J.N., J.P. Kennelly, S. Wan, J.E. Vance, D.E. Vance, and R.L. Jacobs. 2017. The critical role of phosphatidylcholine and phosphatidylethanolamine metabolism in health and disease. BBA - Biomembr. 1859:1558–1572.

27. Lorent, J.H., K.R. Levental, L. Ganesan, G. Rivera-Longsworth, E. Sezgin, M. Doktorova, E. Lyman, and I. Levental. 2020. Plasma membranes are asymmetric in lipid unsaturation, packing and protein shape. Nat. Chem. Biol. 16:644–652.

28. Brennan, P.J., and H. Nikaido. 1995. The Envelope of Mycobacteria. Annu. Rev. Biochem. 64:29–63.

29. Marsh, D. 2013. Handbook of Lipid Bilayers. 2nd ed. CRC Press, Boca Raton.

30. Verkleij, A.J., R.F.A. Zwaal, B. Roelofsen, P. Comfurius, D. Kastelijn, and L.L.M. van Deenen. 1973. The asymmetric distribution of phospholipids in the human red cell membrane. A combined study using phospholipases and freeze-etch electron microscopy. BBA - Biomembr. 323:178–193.

31. Yang, S.-T., A.J.B. Kreutzberger, J. Lee, V. Kiessling, and L.K. Tamm. 2016. The role of cholesterol in membrane fusion. Chem. Phys. Lipids. 199:136–143.

32. Dietrich, C., L.A. Bagatolli, Z.N. Volovyk, N.L. Thompson, M. Levi, K. Jacobson, and E. Gratton. 2001. Lipid Rafts Reconstituted in Model Membranes. Biophys. J. 80:1417–1428.

33. Veatch, S.L. 2007. Electro-formation and fluorescence microscopy of giant vesicles with coexisting liquid phases. Methods Mol. Biol. 398:59–72.

34. Pautot, S., B.J. Frisken, and D.A. Weitz. 2003. Production of Unilamellar Vesicles Using an Inverted Emulsion. Langmuir. 19:2870–2879.

35. Cooper, A., and A.B. Subramaniam. 2024. Ultrahigh yields of giant vesicles obtained through mesophase evolution and breakup. bioRxiv, doi: 10.1101/2024.06.03.597257 (preprint posted June 4, 2024).

36. Kim, E., O. Graceffa, R. Broweleit, A. Ladha, A. Boies, and R.J. Rawle. 2024. Lipid loss and compositional change during preparation of liposomes by common biophysical methods. bioRxiv, doi: 10.1101/2024.05.30.596670 (preprint posted June 2, 2024).

37. Grusky, D.S., A. Bhattacharya, and S.G. Boxer. 2023. Secondary Ion Mass Spectrometry of Single Giant Unilamellar Vesicles Reveals Compositional Variability. J. Am. Chem. Soc. 145:27521–27530.

38. Koynova, R., and M. Caffrey. 1998. Phases and phase transitions of the phosphatidylcholines. BBA - Rev. Biomembr. 1376:91–145.

39. Silvius, J.R., and R. N. McElhaney. 1979. Effects of Phospholipid Acyl Chain Structure on Thermodynamic Phase Properties. 2: Phosphatidylcholines with Unsaturated or Cyclopropane Acyl Chains. Chem. Phys. Lipids. 25:125–134.

40. Findlay, E.J., and P.G. Barton. 1978. Phase behavior of synthetic phosphatidylglycerols and binary mixtures with phosphatidylcholines in the presence and absence of calcium ions. Biochemistry. 17:2400–2405.

41. Koynova, R., and M. Caffrey. 1994. Phases and phase transitions of the hydrated phosphatidylethanolamines. Chem. Phys. Lipids. 69:1–34.

42. Witkowska, A., L. Jablonski, and R. Jahn. 2018. A convenient protocol for generating giant unilamellar vesicles containing SNARE proteins using electroformation. Sci. Rep. 8:9422.

43. Moga, A., N. Yandrapalli, R. Dimova, and T. Robinson. 2019. Optimization of the Inverted Emulsion Method for High-Yield Production of Biomimetic Giant Unilamellar Vesicles. ChemBioChem. 20:2674–2682.

44. Bligh, E.G., and W.J. Dyer. 1959. A rapid method of total lipid extraction and purification. Can. J. Biochem. Physiol. 37:911–917.

45. Hines, K.M., J.C. May, J.A. McLean, and L. Xu. 2016. Evaluation of Collision Cross Section Calibrants for Structural Analysis of Lipids by Traveling Wave Ion Mobility-Mass Spectrometry. Anal. Chem. 88:7329–7336.

46. Hines, K.M., J. Herron, and L. Xu. 2017. Assessment of altered lipid homeostasis by HILIC-ion mobility-mass spectrometry-based lipidomics. J. Lipid Res. 58:809–819.

47. Li, A., K.M. Hines, and L. Xu. 2020. Lipidomics by HILIC-Ion Mobility-Mass Spectrometry. Methods Mol. Biol. 2084:119–132.

48. Ross, D.H., J.H. Cho, R. Zhang, K.M. Hines, and L. Xu. 2020. LiPydomics: A Python Package for Comprehensive Prediction of Lipid Collision Cross Sections and Retention Times and Analysis of Ion Mobility-Mass Spectrometry-Based Lipidomics Data. Anal. Chem. 92:14967–14975.

49. Herron, J., K.M. Hines, and L. Xu. 2018. Assessment of altered cholesterol homeostasis by xenobiotics using ultra-high performance liquid chromatography-tandem mass spectrometry. Curr. Protoc. Toxicol. 78:e65.

50. Garg, S., F. Castro-Roman, L. Porcar, P. Butler, P.J. Bautista, N. Krzyzanowski, and U. Perez-Salas. 2014. Cholesterol solubility limit in lipid membranes probed by small angle neutron scattering and MD simulations. Soft Matter. 10:9313–9317.

51. Shaw, T.R., K.C. Wisser, T.A. Schaffner, A.D. Gaffney, B.B. Machta, and S.L. Veatch. 2023. Chemical potential measurements constrain models of cholesterol-phosphatidylcholine interactions. Biophys. J. 122:1105–1117.

52. Cornell, C.E., A. Mileant, N. Thakkar, K.K. Lee, and S.L. Keller. 2020. Direct imaging of liquid domains in membranes by cryo-electron tomography. Proc. Natl. Acad. Sci. 117:19713–19719.

53. Heberle, F.A., M. Doktorova, H.L. Scott, A.D. Skinkle, M.N. Waxham, and I. Levental. 2020. Direct label-free imaging of nanodomains in biomimetic and biological membranes by cryogenic electron microscopy. Proc. Natl. Acad. Sci. 117:19943–19952.

54. Heberle, F.A., M. Doktorova, S.L. Goh, R.F. Standaert, J. Katsaras, and G.W. Feigenson. 2013. Hybrid and Nonhybrid Lipids Exert Common Effects on Membrane Raft Size and Morphology. J. Am. Chem. Soc. 135:14932–14935.

55. Morita, M., H. Onoe, M. Yanagisawa, H. Ito, M. Ichikawa, K. Fujiwara, H. Saito, and M. Takinoue. 2015. Droplet-Shooting and Size-Filtration (DSSF) Method for Synthesis of Cell-Sized Liposomes with Controlled Lipid Compositions. ChemBioChem. 16:2029–2035.

56. Dimitrov, D.S., and M.I. Angelova. 1988. Lipid swelling and liposome formation mediated by electric fields. Bioelectrochem. Bioenerg. 19:323–336.

57. Pazzi, J., and A.B. Subramaniam. 2020. Nanoscale Curvature Promotes High Yield Spontaneous Formation of Cell-Mimetic Giant Vesicles on Nanocellulose Paper. ACS Appl. Mater. Interfaces. 12:56549–56561.

58. Horger, K.S., D.J. Estes, R. Capone, and M. Mayer. 2009. Films of Agarose Enable Rapid Formation of Giant Liposomes in Solutions of Physiologic Ionic Strength. J. Am. Chem. Soc. 131:1810–1819.

59. A. Weinberger, A. F.-C. Tsai, G.H. Koenderink, T.F. Schmidt, R. Itri, W. Meier, T. Schmatko, Schröder, and C. Marques. 2013. Gel-Assisted Formation of Giant Unilamellar Vesicles. Biophys. J. 105:154–164.

59. MacDonald, R.C., R.I. MacDonald, B.Ph.M. Menco, K. Takeshita, N.K. Subbarao, and L. Hu. 1991. Small-volume extrusion apparatus for preparation of large, unilamellar vesicles. Biochim. Biophys. Acta BBA - Biomembr. 1061:297–303.

